# Substrate-binding Glycine Residues are Major Determinants for Hydrolase and Ligase Activity of Plant Legumains

**DOI:** 10.1101/2022.09.26.509423

**Authors:** Xinya Hemu, Ning-Yu Chan, Heng Tai Liew, Side Hu, Xiaohong Zhang, Aida Serra, Julien Lescar, Chuan-Fa Liu, James P Tam

**Author notes:** **Corresponding Author** James P. Tam.

## Abstract

Peptide asparaginyl ligases (PALs) are Asn/Asp(Asx)-specific ligases that are useful for precision modifications of proteins and live-cell surfaces. However, PALs share high structural similarity to the far more common asparaginyl endopeptidases (AEPs), also known as legumains that hydrolyze peptide bonds after Asx, thus making it challenging to identify PALs in a sea of AEPs. Previously we identified sequences flanking the catalytic site as ligase activity determinants (LADs) for legumains. Here we show that two conserved substrate-binding Gly residues are critical, but negative determinants for ligase activity, based on a combined bioinformatics analysis of 1,500 plant legumains, mutagenesis and functional study of 16 novel legumains, plus identification of seven new PALs. We also show that PALs are rare and AEPs are much more common, accounting for about 1% and 88%, respectively. Our results suggest that specific glycine residues are molecular determinants to identify PALs and AEPs as two different legumain subfamilies.

## INTRODUCTION

Peptide ligases form peptide bonds whereas proteases break them. Ligases are rare in nature, but can be found as bio-processors of ribosomally-synthesized and post-translationally modified peptides (RiPPs) in bacteria,(Mazmanian et al., 1999) cyanobacteria,(Lee et al., 2009) fungi,(Luo et al., 2014) and plants.(Barber et al., 2013, Nguyen et al., 2014) In particular, plants produce peptide asparaginyl ligases (PALs) for biosynthesis of N-to-C cyclic peptides, forming an Asn/Asp (Asx) peptide bond with N-terminal residues.

Together with asparaginyl endopeptidases (AEPs), PALs belong to Cys proteases of the C13 subfamily (MEROPS EC 3.4.22.34).(Rawlings et al., 2018) They are also known as vacuolar processing enzymes (VPEs) because they function in lytic vacuoles as degradative enzymes,(Hara-Nishimura et al., 1991) or legumains because they were discovered in legumes.(Kembhavi et al., 1993, Takeda et al., 1994) Legumains are expressed as proenzymes in which the core domain is protected by an inhibitory cap domain that can be removed by acid-induced auto-activation. In plants, legumains regulate diverse cellular processes including maturation of defence peptides and proteins, degradation of storage proteins during seed germination, and programmed cell death.(Hara-Nishimura et al., 1995, Hiraiwa et al., 1997, Hatsugai et al., 2004)

Legumains also cut and join single peptide or protein substrates. This legumain-mediated splicing process was first found in maturation of lectins.(D. M. Carrington et al., 1985, Min and Jones, 1994) Recent examples are the biosynthesis of sunflower seed trypsin inhibitors(James et al., 2018) and the cyclic trypsin inhibitor MCoTIs.(Du et al., 2020, Liew et al., 2021) Both biosynthetic reactions involve a cleavage first at an Asn site and then ligation at a downstream Asp to form a cyclic peptide. Although the hydrolase and splicing activity of AEP has been known for over 40 years, the Asx-specific ligase activity of a PAL was only confirmed in 2014 with the discovery of butelase-1 as the first PAL.(Nguyen et al., 2014) Butelase-1 displays efficient ligase activity under both acidic and neutral pH. In contrast to the more common hydrolytic legumains, butelase-1 has almost no protease activity. Subsequent identification of additional butelase-1-like ligases(Harris et al., 2015, Harris et al., 2019, Jackson et al., 2018, Hemu et al., 2019a) firmly established PALs as ligating legumains that are functionally distinct from both hydrolytic legumains (AEPs) and the splicing legumains.

PALs are versatile and precise tools for site-specific modification and can catalyze semi-and totalsyntheses of peptides and proteins. PAL-mediated modifications include protein cyclization,(Nguyen et al., 2015b, Hemu et al., 2016, Nguyen et al., 2016) live-cell labeling,(Bi et al., 2017, Bi et al., 2020) conjugation,(Nguyen et al., 2015a, Bi et al., 2018) and polymerization.(Cao et al., 2016, Hemu et al., 2019b) Several chemical and enzymatic ligation methods for bio-orthogonal or chemoenzymatic ligation either in tandem or under one-pot conditions are also compatible with PALs.(Cao et al., 2015, Harmand et al., 2018, Wang et al., 2021)

Attempts to find a structural basis to distinguish PAL from AEPs have largely been unsuccessful. Comparison of 12 unique crystal structures of AEP and PAL core domains gave a RMSD of <1 Å.(Hemu et al., 2020) Recently, we showed that certain sequences flanking the catalytic S1 cysteine pocket could serve as ligase-activity determinants (LADs) to distinguish a PAL from an AEP.(Hemu et al., 2019a, Hemu et al., 2020) Here, we predicted and then experimentally validated PALs as a distinct subfamily of legumains in an analysis that refined LAD hypothesis based on bioinformatics study of 1,500 plant legumains.

## MATERIALS AND METHODS

### Retrieval of plant legumain sequences from existing protein and transcriptome databases

Core domain sequences of butelase-1 (from G44 to N324) (KF918345), butelase-2 (ALL55651.1), OaAEP1b (KR259377), AtVPE-α (NP_180165.1), AtVPE-β (NP_176458.1), and AtVPE-γ (NP_195020.1) were used as queries to mine transcriptomes deposited in NCBI and OneKP (Matasci et al., 2014) databases using BLASTp (nr) and tBLASTx (nr and TSA) searches. The selection threshold were set to >50% identity and >95% sequence coverage as compared to the queries. Data from all searches were combined, and repeated homologs were removed to subsequently retrieve 1,498 unique ORFs that encode plant legumains including 495 sequences from OneKP and 1,003 sequences from NCBI. Two previously reported PALs, OaAEP4 and OaAEP5, were not found in the transcriptomic database and were added manually (Supplementary Dataset 1).

#### Bioinformatics analysis

Multiple sequence alignments were performed using Clustal Omega with BLOSUM62 substitution matrix and default gap penalties (Sievers et al., 2011). Aligned sequences were submitted to the UET server (http://lichtargelab.org/software/uet) for Universal evolutionary trace analysis to find functionally-important residues. The rvET rank scores were mapped to the butelase-1 crystal structure (PDB code: 6DHI) and visualized using PyMOL. The phylogenetic tree built from the UET analysis was uploaded to iTOL for annotation. Sequence logos were extracted from Jalview 2.10.

#### Cloning, expression and purification of recombinant proteins

cDNA sequences of 16 selected enzymes (VyPAL4, MK085233.1, VyPAL5, MK085234.1; VaPAL1, GFWC01037197.1; VoPAL1, GFXR01024405.1; VuPAL1, GCAB01004088.1, VvPAL1, GFWF01025417.1; HaPAL1, LOC110864271; BmAEP1, XM_022292359.1; PiAEP1, OneKP:BQEQ_2002574; LsAEP1, JI582927.1; PeAEP1, GBRT01052954.1; PeAEP2, GBRT01019050.1; DcAEP1, OneKP:SHEZ_2097111; BrAEP1, GFUS01046649.1; CrAEP1, GACD01060809.1; VtAEP1, OneKP:LPGY_2014619) were codon-optimized for E. coli expression system. Sequences starting after the residue corresponding to butelase-1-L26 or after the signal peptide predicted using SignalP5.0 were cloned into the pET28a(+) vector at Ndel/Xhol restriction sites to generate a His6-fusion protein construct (Genscript, USA). Point mutations were generated using a Q5 mutagenesis kit (New England Biolabs, USA). Plasmids were amplified in DH5α E. coli and protein was expressed from SHuffle T7 E. coli competent cells (S3026J, New England Biolabs, USA). The transformed cells were grown to OD600 = 0.4 at 30 °C in LB broth supplemented with 50 μg/mL kanamycin. The temperature was reduced to 16 °C and expression was induced by 0.1 mM isopropyl β-D-1-thiogalactopyranoside for 24-48 h. Cells were harvested by centrifugation at 6,000 g for 15 min at 4 °C and the pellet was re-suspended in cold lysis buffer (20 mM Na HEPES, 100 mM NaCl, 1mM EDTA, 1 mM dithiothreitol (DTT), and 0.1 % (v/v) Triton X-100, pH 7.0). Bacterial cells were lysed by sonication on ice and cell debris was removed by centrifugation at 10,000 g for 30 min at 4 °C. The filtered supernatant was loaded onto a column packed with cOmpleteTM His-tag purification resin (Merck). His6-proteins were eluted with 4x 2 bead volumes of elution buffer (20 mM Na HEPES, 1 mM EDTA, 1 mM DTT, 0.5 M imidazole, pH 7.5). Eluents were concentrated and buffer-exchanged with binding buffer (20 mM Na HEPES, 1 mM EDTA, 1 mM DTT, pH 7.5) using a centrifugal filter unit (Vivaspin-15, Satorius) to remove excessive imidazole. Additional purification was carried out by anion-exchange chromatography on a 1 mL HiTrap Q Sepharose column (GE Healthcare, USA). Yields of the recombinant-produced proenzymes ranged from 0.1 to 2 mg per litre culture. Active butelase-2 used in this study was prepared as previously described using a Baculovirus system and protein expression in insect cells (Hemu et al., 2020).

#### Acid-induced auto-activation and purification of active enzymes

Purified proenzymes in a concentration of 1-2 mg/mL were activated by lowering the pH to 4.0-4.5 and incubating at 37 °C for 10 min to 2 h. For VaPAL-I342A and HaPAL1-V345A, activation was performed at pH 4.0-4.1 at 25 °C overnight. Activation progress was analyzed by SDS-PAGE. Activated enzymes were purified on a HiLoad 16/600 Superdex 75 column with 20 mM sodium citrate buffer, 1 mM EDTA, 1 mM DTT, 100 mM NaCl and 5% (w/v) glycerol, pH 4.2. The concentration of purified enzymes was determined by measuring the absorbance at 280 nm on a NanoDrop 2000 spectrophotometer (Thermo Fisher, USA). The eluents were neutralized to pH 5-6 and stored at 4 °C or −80 °C after the addition of 20% sucrose.

#### Determination of auto-activation sites

After separation by SDS-PAGE, the bands for activated enzymes were excised from the gels and subjected to in-gel tryptic digestion. Digested peptide fragments were extracted and sequenced by LC-MSMS as previously described (Serra et al., 2016).

#### Functional studies

Intramolecular cyclization of peptide substrates were performed at 37 °C in reaction buffers having pH ranging from 4 to 8 (20 mM citrate or phosphate buffers with 1 mM EDTA and 5 mM β-mercaptoethanol). The peptide substrates GN12-GL (GLYRRGRLYRRN-GL) and SFTI1-GL (GRCTKSIPPICFPD-GL) were synthesized using Fmoc chemistry on automated synthesizer (LibertyBlue, CEM) and purified by preparative RP-HPLC. Activated recombinant legumains and substrates were mixed to the final concentrations of 40 nM and 20 μM, respectively. The reactions were monitored by MALDI-TOF mass spectrometry (5800 Applied Biosystem, USA) and quantitatively analyzed by RP-HPLC on a C18 analytical column (AerisTM widepore, Phenomenex, USA) after being quenched with 1:1 v/v acetonitrile with 0.1% trifluoroacetic acid.

## RESULTS

### LADs based on substrate-binding residues at P and P’ sites

Currently, eight PALs have been identified based on mutagenesis and functional studies. Sequence comparison and structural analysis of the reported PALs and AEPs suggested two LAD sites, LAD1 and 2, correspond to the substrate-binding residues (Fig. 1).(Zauner et al., 2018, Hemu et al., 2020, Chen et al., 2021) LAD1, located at the non-prime, or amino-side, of the Asx-Xaa scissile peptide bond, is a hydrophobic tripeptide forming the βIV strand that shapes the S2 substrate-binding pocket.(Aaslanda et al., 2002) Mutagenesis studies suggested that the middle residue of the LAD1 tripeptide, which in butelase-1 corresponds to Val237, and that is also known as the “gate-keeper(Yang et al., 2017), exerts a stronger influence on enzymatic directionality than the first and the third residue. At this middle position, all eight known PALs possess a hydrophobic or bulky residue, including Val, Ile, Cys, and Pro.(Hemu et al., 2020) In contrast, known protease-legumains, or AEPs, such as AtVPEs, HaAEP1 and butelase-2, contain a Gly residue in the middle position of the LAD1 motif.

**Figure 1.**
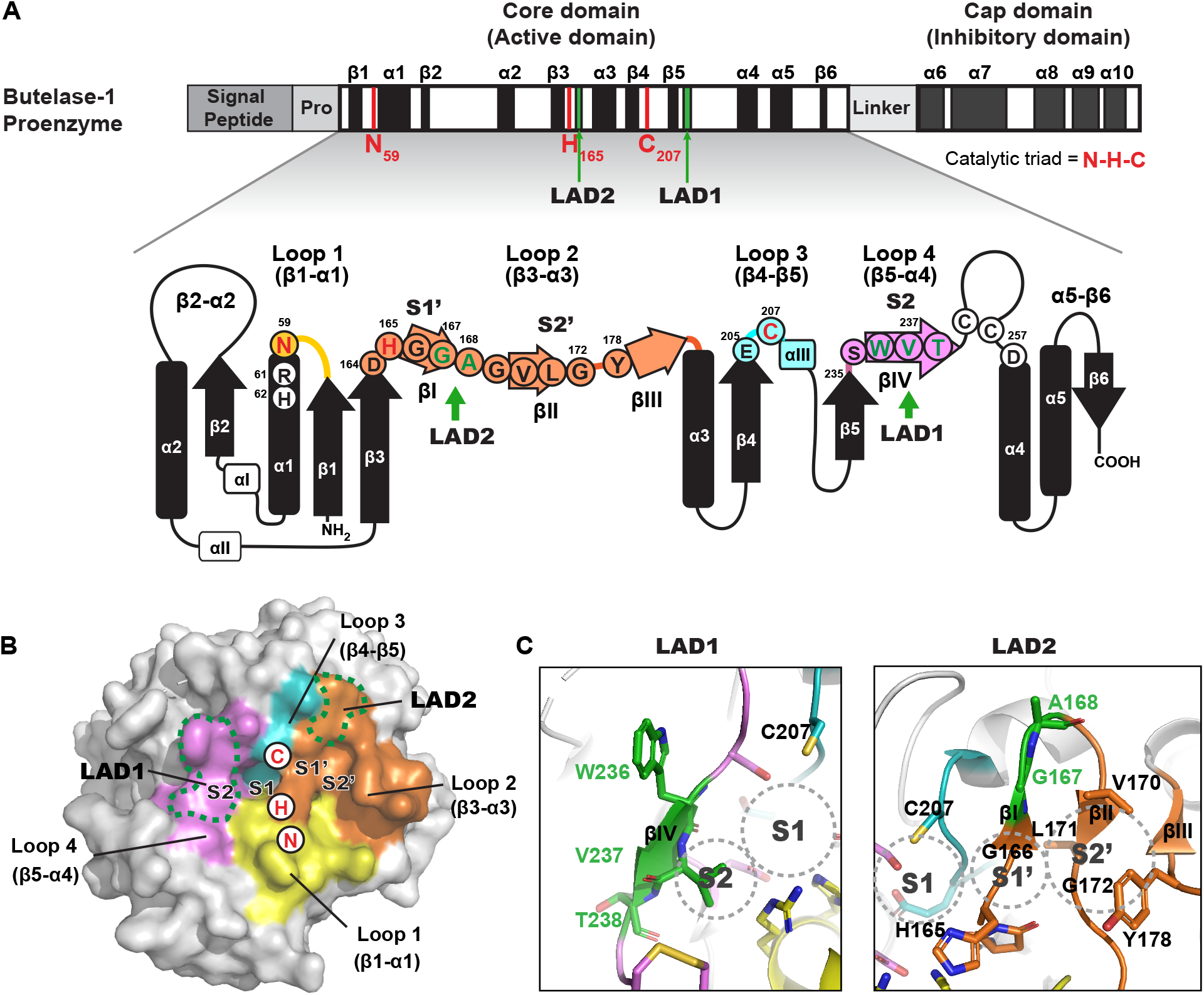
1D, 2D, and 3D illustration of the ligase-activity determinants LAD1 and LAD2 on butelase-1. (A) 1D schematic and 2D topology diagram of butelase-1. Location of LAD motifs in substrate-binding loop 2 and 4 on top of the core domain are labeled. Residues forming substrate-binding pockets S2-S1-S1’-S2’ are circled. Catalytic residues His165 and Cys207 are colored in red. LAD residues are colored in green. Other residues are in blank. (B) Four major substrate-binding loops mapped on the 3D structure of butelase-1 (PDB access code: 6DHI). Loop1-4 are colored in yellow, orange, cyan, and magenta, respectively. LAD motifs are marked with green dashed lines on the molecular surface near the S1 oxyanion hole. (C) Close-up view of LAD1 tripeptide W236-V237-T238 that form the S2 pocket and LAD2 dipeptide G167-A168 next to S1’ and S2’ pockets.

LAD2, located at the prime, or carboxyl side, of the scissile bond, is an aliphatic tripeptide corresponding to a ligating-legumain such as butelase-1 Gly166-Gly167-Ala168 that overlaps with the βI strand. Because Gly166 in the S1’ pocket is conserved in AEPs and PALs, LAD2 was reduced to a dipeptide motif. Examples of PAL-like LAD2 motifs include Gly167-Ala168 (butelase-1), Ala-Pro (OaAEP3/4, HeAEP3, VyPAL1/2) and Ala-Ala (OaAEP1b, OaAEP5). In contrast, most known AEPs have Gly167-Pro168.

### Data collection for 1,500 plant legumains

To confirm the current LADs using a large data set, we collected plant legumain sequences from existing databases by BLAST search of the National Centre for Biotechnology Information (NCBI) and one-thousand-plants (OneKP) database using core-domain protein sequences (Gly44-Asn324, butelase-1 numbering) of known PALs and AEPs as queries.(Matasci et al., 2014) Sequences with <50% identity were excluded because most belong to GPI-anchoring transamidases, another type of C13 Cys protease.(Ohishi et al., 2000) Together, we obtained 1,500 non-redundant sequences including 1,003 sequences from NCBI, 495 sequences from OneKP, and two additional PAL sequences, OaAEP4 and OaAEP5.(Harris et al., 2019) (supplement Dataset 1)

The 1,500 legumain sequences belong to 875 species from 249 plant families, ranging from primitive plants, such as green algae, moss, fern, and conifer, to flowering plants (Supplementary Fig. 1). Crops of economic importance are highly represented in our dataset, including 13% (194/1500) from 75 species of the Poaceae family and 9% (132/1500) from 87 species of the Fabaceae family. There are 50 families containing between 5 and 80 sequences and the remaining 197 families contain <5 sequences. The six plant families that known for producing legumain-processed cyclic peptides, including Rubiaceae, Violaceae, Fabaceae, Solanaceae, Cucurbitaceae, and Asteraceae, together contain 370 sequences from 159 species.

### Evolutionary trace (UET) analysis to reveal evolutionarily-important residues for legumains

To identify molecular determinants for a PAL or an AEP using a different and global method to corroborate our proposed LADs, we performed universal evolutionary trace (UET)(Lua et al., 2016) analysis with the 1,500 aligned sequences to determine the evolutionary importance of each residue. UET analysis gives each residue a real-value Evolutionary Trace (rvET) score(Mihalek et al., 2004) that is calculated based on the level of diversity of a particular residue among distant evolutionary sequences (Supplementary file 2). An rvET score of 1.0 indicates a completely conserved residue that has a critical function. In contrast, a high rvET score of up to 200 indicates the presence of diverse residues among evolutionarily-close analogs that have low functional importance. Fig. 2A shows the rvET score of each residue mapped onto the butelase-1 zymogen crystal structure (PDB ID: 6DHI)(Yang et al., 2017) using a color gradient ranging from red (most important) to white (least important) for visualization.

**Figure 2.**
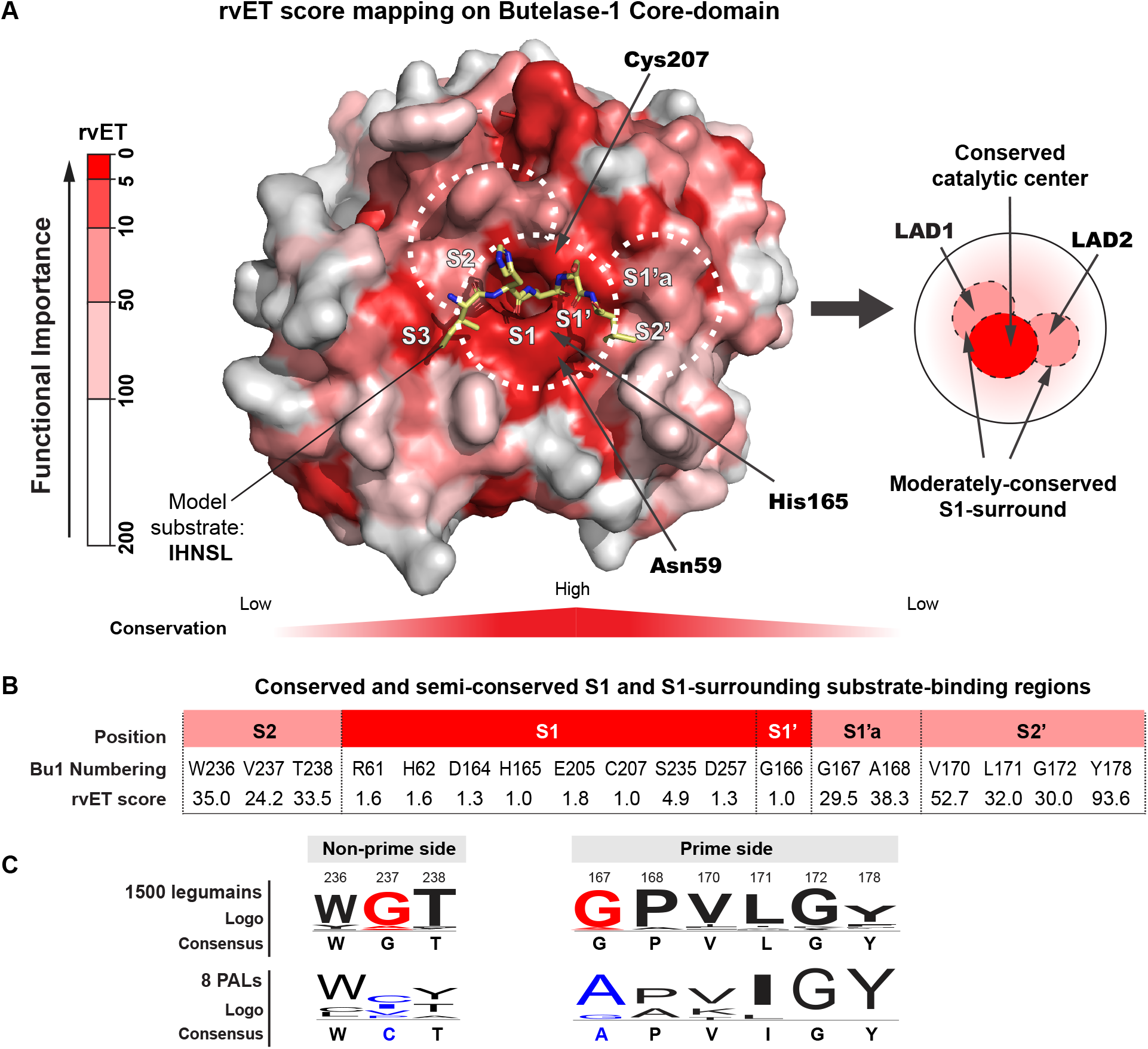
Identification of evolutionarily-important residues of plant legumains by UET analysis. (A) Ribbon structure of butelase-1 (PDB code: 6DHI) colored according to rvET scores. Red and white refer to the most and least conserved positions, respectively. A model peptide substrate IHNSL (yellow) was placed on top of the surface to show the location of substrate binding pockets. An rvET-mapped illustration highlights the conserved central ring (red) and two semi-conserved second rings (salmon). LAD motifs are located in the semi-conserved rings. (B) rvET scores of the moderately-conserved residues. Residues forming S1-S1’ pocket and catalytic dyad are highly conserved with rvET score <5 (red). Away from the catalytic center, the conservation gradually decreased with an increasing of rvET score in the range of 24.2-93.6. (C) Comparison of sequence logos of moderately-conserved S1-surrounding residues between 1500 legumains and 8 confirmed PALs. Conserved Gly237 and Gly167 (red) in plant legumains are occupied by non-Gly residues in PALs (blue).

The substrate-binding sites S1 and S1’ comprise evolutionarily-important residues that have low rvET scores <2 with surrounding residues having rvET scores of around 30 (residues colored red and pink, respectively, in Fig. 2B). This result is consistent with the fact that both AEPs and PALs share the same S1-pocket specificity towards Asx, but have substrate-binding pockets that vary in their enzymatic directionality and specificity.(Dall and Brandstetter, 2013, Zauner et al., 2018)

Further analysis of consensus sequences in the substrate-binding sites of the 1,500 legumains show that the moderately conserved residues are Gly167/Pro168 next to S1’ pocket, Leu171/Gly172 in S2’ pockets, and Trp236/Gly237/Thr238 in S2 pocket. Importantly, two moderately-conserved Gly residues, Gly167 and Gly237, are found to be non-Gly in the consensus sequences of PALs, supporting their importance as the key residues in LAD1 and LAD2 that influence ligase and protease-activity of legumains (Fig. 2C). Overall, the global sequence analysis by UET agrees with the previously proposed LAD motifs.

#### Occurrence and Distribution of PAL-like LADs in legumains

We performed detailed analyses of amino acid occurrences at residue 237 of LAD1 and residue 167 of LAD2, which are occupied by Gly in 90% and 88% of legumains, respectively. Besides Gly, 8% of legumains possess small residues like Ala and Ser at position 237. The remaining 2% (29/1,500) possess a PAL-like bulky amino acid that are grouped as LAD1+ motifs (Fig. 3A). On the prime side, Gly167 followed by a Pro at position 168 was the most common combination and was found in 84% of legumain sequences. Meanwhile, 11% of legumains (166/1,500) contained PAL-like LAD2 dipeptides (Ala167-Xaa168 or Gly167-Ala168), including 92 sequences with an Ala167-Xaa168 (Xaa = Pro, Ala, Thr, Val, Ser) dipeptide and 74 sequences with a butelase-1-like Gly167-Ala168 dipeptide, Which are grouped as LAD2+ motifs. The remaining 5% of sequences had other residues at position 167 including Ser, Tyr, Thr, Asp, Asn, and Val (Table S1).

**Figure 3.**
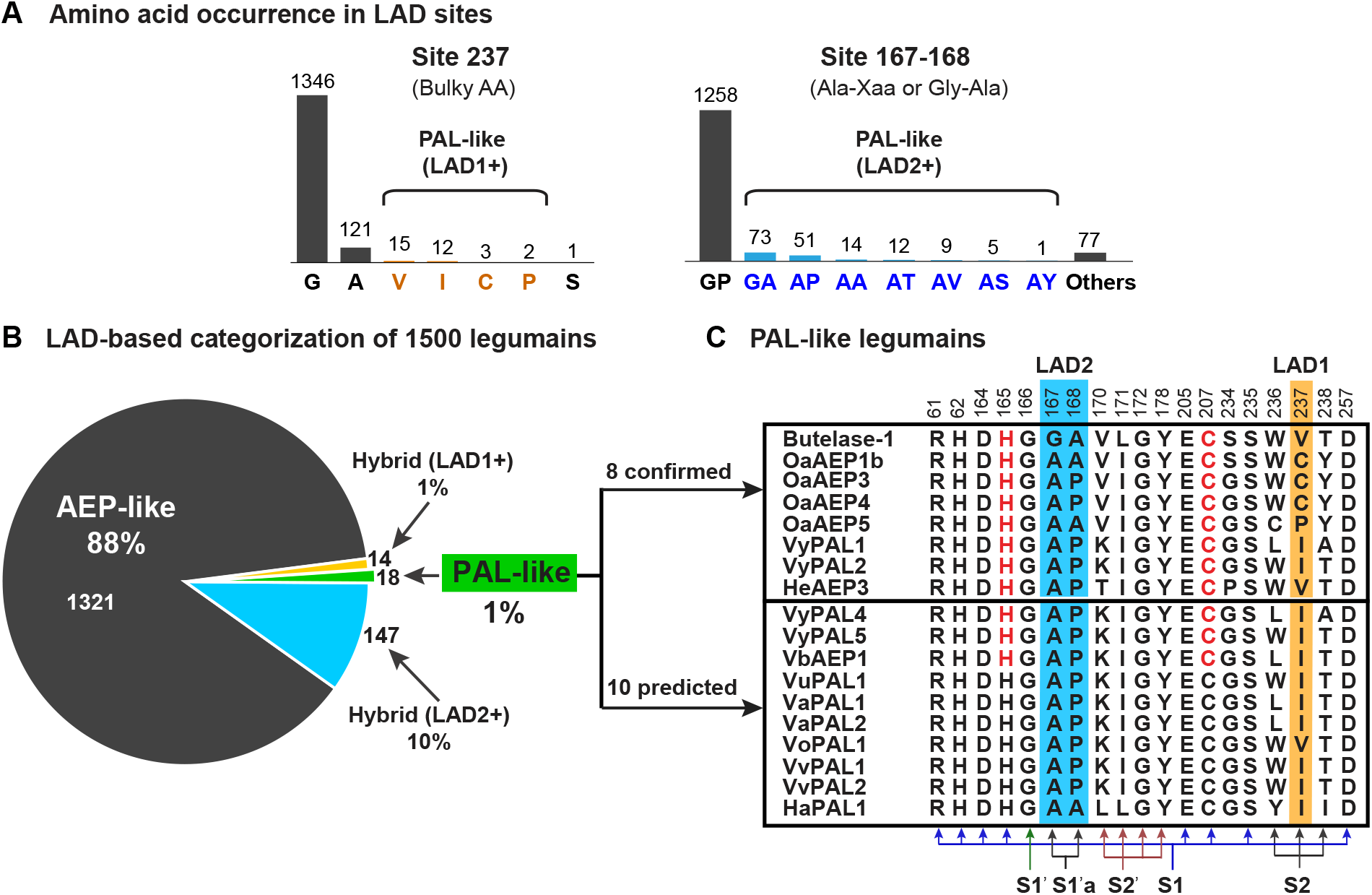
Screening of new PALs based on amino acid composition of LADs. (A) Amino acid occurrences at position 237 (LAD1) and 167-168 (LAD2) of plant legumains. At position 237, Val, Ile, Cys and Pro are predicted to be PAL-like LAD1 motif (LAD1+). At position 167-168, Gly-Ala and Ala-Xaa are predicted to be PAL-like LAD2 motif (LAD2+). The rest are predicted to be AEP-like (LAD-). (B) Categorization of 1500 plant legumains into AEP-like, PAL-like and two Hybrid subtypes based on LAD compositions. 18 sequences are classified as PAL-like legumains with both PAL-like LAD1 and LAD2 motifs. They include all 8 confirmed PALs and 10 predicted PALs as listed in the right panel. Catalytic residues, pocket-forming residues, and LAD sites are aligned and labeled with butelase-1 numbering.

Based on the amino acid composition at position 237 and 167-168, we grouped the 1,500 legumains into three types: AEP-like (hydrolytic-), PAL-like (ligating-) and hybrid legumains (Fig. 3B). AEP-like legumains, that lack LAD+ motifs, are most common, and present in 88% of the 1,500 legumains. In contrast, only 18 sequences correspond to PAL-like legumains, accounting for only 1.1% of the sequences. These include 8 functionally-characterized PALs, and 10 predicted PALs that are named after their host species Viola albida, V. betonicifolia, V. orientalis, V. uliginosa, V. verecunda, V. yedoensis and Helianthus annuus (Fig. 3C). Notably, the newly-predicted PAL-like legumains are all derived from plants that produce cyclic peptides. Residue 237 of LAD1 in these PAL-like sequences is occupied by bulky and hydrophobic amino acids, whereas the LAD2 motif (residues 167-168) has either an Ala-Pro or Ala-Ala dipeptide sequence. The remaining 10% of legumains are hybrids, with 14 sequences bearing a LAD1+ motif and 148 sequences bearing a LAD2+ motif.

### Functional validation of LAD-based predictions

To validate the LAD-based classification, we selected 15 predictions for recombinant expression and activity tests. This group included 7 PALs, 3 AEPs and 5 hybrids. For comparison, we also included the previously characterized butelase-2 (Nguyen et al., 2014, Serra et al., 2016) as an AEP control.

Proenzymes of each legumain were expressed in E. coli using a protocol we previously described41. Acid-induced auto-activation at pH 4.0-4.5 was successful for all except VaPAL1 and HaPAL1 (Supplementary Fig. 2). For these two legumains, we found that substitution of a conserved hydrophobic residue in the linker region, VaPAL1-Ile342 or HaPAL1-V345, with alanine could facilitate auto-activation (Supplementary Fig. 3). This mutation site is located after the major autolytic cleavage sites as confirmed by MS/MS sequencing (Supplementary Fig. 4) and thus will not remain in the activated enzyme.

All functional studies of recombinant legumains were performed with the modified sunflower-trypsin-inhibitor peptide substrate GN12-GL (GLYRRGRLYRRN-GL, M.W. 1,749 Da), which yields the hydrolyzed linear peptide GN12 (M.W. 1579 Da) or the N-to-C cyclized peptide cGN12 (M.W. 1561 Da) through Asn-specific hydrolysis or cyclization, respectively (Fig. 4A). The common recognition signal of Asn-Gly-Leu was chosen based on the known substrate specificity of characterized legumains.(Nguyen et al., 2014, Hemu et al., 2019a, Dall et al., 2020) To show that legumain-mediated catalytic reactions are pH-dependent,(Nonis et al., 2021, Bernath-Levin et al., 2015) we performed functional studies at nine discrete pHs, from pH 4 to 8, with an interval of 0.5 pH unit. A fixed enzyme-to-substrate ratio of 1:500 (mol: mol) was applied and reactions were allowed to proceed at 37 °C for 5 or 10 min. Progress of the reactions was monitored by MALDI-TOF MS and yields were quantified by RP-HPLC (Supplementary Fig. 5).

**Figure 4.**
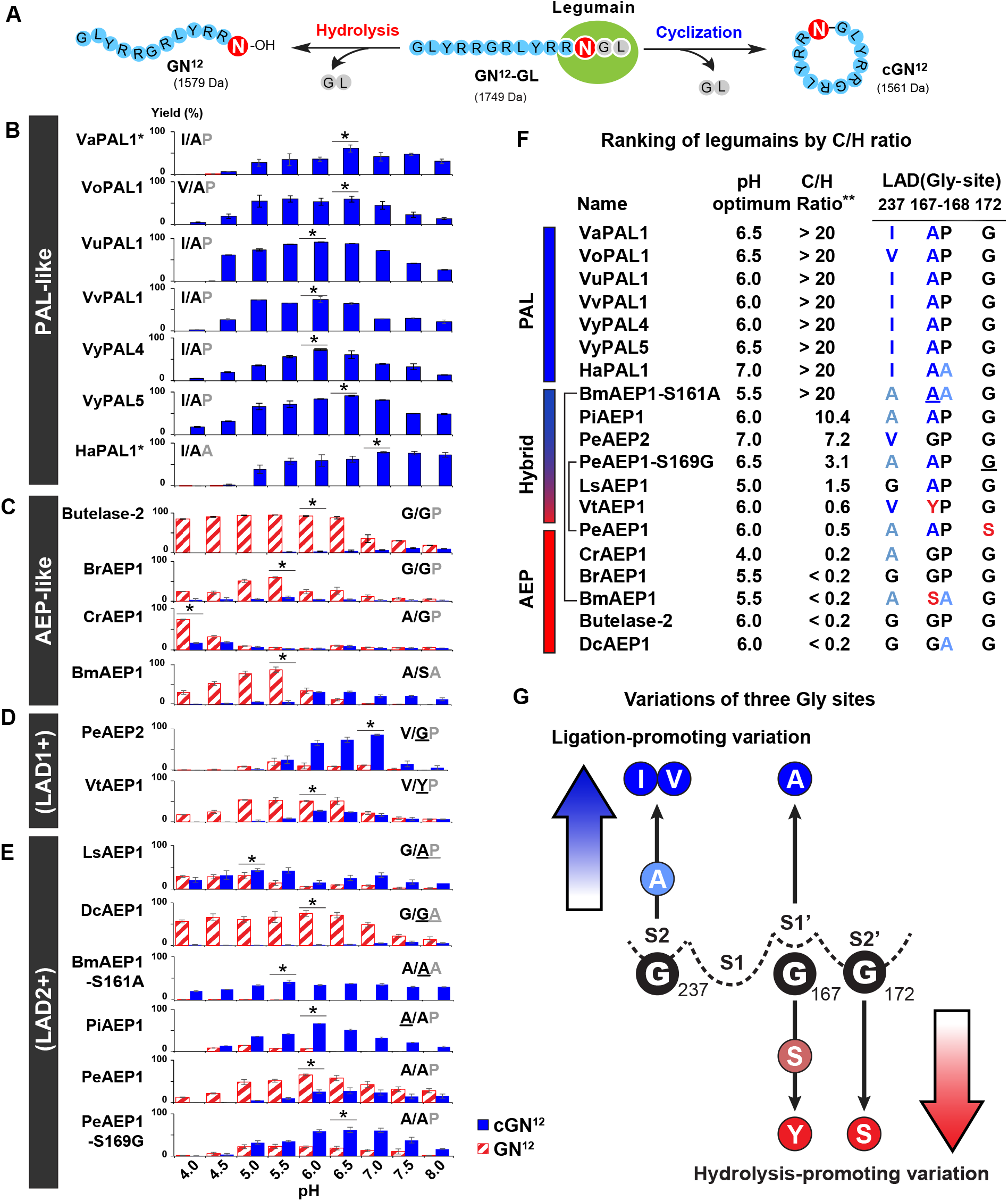
Activity study of selected plant legumains. (A) Legumain-mediated reactions with the model substrate GN12-GL could yield both hydrolytic product GN12 and cyclized product cGN12 with 18 Da mass difference. (B) The product distribution profiles of PAL-like legumains, (C) AEP-like legumains, (D) Hybrid legumains with LAD1+ motifs and (E) Hybrid legumains with LAD2+ motifs. Reactions were performed with a fixed enzyme-to-substrate molar ratio of 1:500 at 37 °C under nine different pHs (pH 4.0 to 8.0) for 5 or 10 min. The product yields (%) were quantified by analytical RP-HPLC (see Figure S8 for raw MS and HPLC data). Mean value and error bar were calculated based on three to five independent reactions. LAD composition of each enzyme are marked in the top corner of the plot. Mutation sites are underlined. Optimal pH for each enzyme is marked with *. (F) Ranking of C/H ratios of 19 recombinant legumains showed over 100-fold difference between PALs and AEPs. **: C/H ratio >20 was an estimated value indicating that no hydrolytic products were detected by HPLC, and so the hydrolysis yield was set as <4% based on the detection limit of HPLC. See full data with statistical analysis in supplementary Table S2. (G) Directionality of legumains are primarily determined by amino acid variations at three conserved and substrate-binding Gly sites.

The seven selected PAL-like legumains (LAD1+ and LAD2+) are VaPAL1, VoPAL1, VuPAL1, VvPAL1, HaPAL1, VyPAL4 and VyPAL5. Two other predicted PALs, VaPAL2 and VvPAL2, were not selected because they are isoforms that share 96.1% and 98.9% sequence identity with VaPAL1 and VvPAL1, respectively. VbAEP1 was previously reported as the only PAL present in transcriptome analyses of its host species.(Rajendran et al., 2021) Functional studies showed that all seven predictions indeed acted as PALs, affording high ligation selectivity with a cyclization/hydrolysis (C/H) ratio >20, a ratio suggesting the products contain <4% GN12 from hydrolysis and >96% cGN12 from ligation. This selectivity was maintained from pH 4 to 8 with no observable hydrolysis at ≥pH 6 (Fig. 4B, Supplementary Table 2). Similar to butelase-1, six predicted Viola PALs displayed pH optima between pH 6 and 6.5. HaPAL1 was the exception with a pH optimum of 7.0-7.5. At pH 6, HaPAL1 could accept both Asp and Asn when tested with a substrate based on the native linear precursor of the sunflower trypsin inhibitor SFTI-GL (GRCTKSIPPICFPD-GL, M.W. 1703 Da) with P1-Asp (Supplementary Fig. 6) to give >90% cyclic SFTI-1, suggesting that it could be a cyclase for SFTI1 bioprocessing. Overall, LAD+ motifs work well in predicting PALs. Among the seven new PALs, VyPAL5 and VuPAL1 are the most efficient, affording >90% cyclization yield within 10 min, whereas the rest gave 60-80% yield.

Among four predicted AEPs (LAD1- and LAD2-), butelase-2 and BrAEP1 (Brassica chinensis) contain the most common LAD combinations of Gly237/Gly167-Pro168. This combination is found in 81% (1213/1500) of legumains in our data set. CrAEP1 (Catharanthus roseus) and BmAEP1 (Momordica charantia, bitter melon) have an Ala-substitution at position 237, which is the second most common amino acid at this position (8%). BmAEP1 has an additional Ser-substitution at position 167, which is the third most commonly found residue with 3.4% occurrence, after Gly (90%) and Ala (6%). For four AEPs, protease activity predominates with C/H ratios <0.25 at their optimal pH, which ranged from pH 4 to 5.5 (Fig. 4C). Similar to butelase-2, all three putative AEPs behaved as predicted.

Since functional studies of hybrid legumains (LAD1+/- and LAD2+/-) could shed light on the relative importance of each LAD, we selected two legumains with LAD1+ and four with LAD2+. LAD1+ hybrids include PeAEP2 (Petunia exserta) and VtAEP1 (Viola tricolor). They have same LAD1-Val237 but different LAD2-motifs as Gly167-Pro168 and Tyr167-Pro-168, respectively. PeAEP2 is bifunctional, acting predominantly as a protease at pH <5, but as a ligase at pH >5.5, with a C/H ratio 7.2 around pH 7 (Fig. 4D). In contrast, VtAEP1, which is a predicted hybrid of LAD1-, acted predominantly like an AEP, with C/H ratios that remained <1 throughout the tested pH range. These results suggest that LAD1+ is not the major determining factor, at least not under acidic conditions of pH <5.5.

Of the four selected LAD2+ hybrids (LsAEP1 from Lactuca sativa, DcAEP1 from Dianthus caryophyllus, PiAEP1 from Psychotria ipecacuanha, and PeAEP1 from Petunia exserta), LsAEP1 and DcAEP1 share the same LAD1-motif of Trp236-Gly237-Thr238, but a different LAD2+ dipeptide motif of Ala167-Pro168 and Gly167-Ala168, respectively. We thus examined which LAD2+ motif confers more PAL-like activity. Functional studies showed that LsAEP1 and DcAEP1 displayed contrasting profiles. LsAEP1 (Ala167-Pro168) had a bifunctional profile with a C/H ratio 1.5 at its optimal pH 5, whereas DcAEP1 (Gly167-Ala168) had a butelase-2-like profile with C/H ratios <0.25 from pH 4 to 8 (Fig. 4E).

To determine the relative importance of Ala167 or Ala168 in LAD2, we performed a LAD2+ Ala-substitution at BmAEP1-Ser161 (corresponding to Ala167 in butelase-1 numbering) to give BmAEP1-S161A (Supplementary Fig. 7). This single mutation efficiently suppressed protease activity of wild-type BmAEP1 and conferred a PAL-like activity with C/H ratios >20 from pH 4.5 to pH 8. These results suggested the importance of LAD2+ and that Ala at position 167 imposed a more profound ligation-promoting effect than Ala at position 168, a result that agrees with the UET analysis.

Two other hybrids, PiAEP1 and PeAEP1, have identical LAD combinations of LAD1- and LAD2+ of Ala237/Ala167-Pro168. However, they displayed very different functional profiles: PiAEP1 was PAL-like (C/H ratio 10.4 at pH 6) and PeAEP1 was AEP-like (C/H ratio 0.49 at pH 6) (Fig. 4F). To explain this 20-fold difference in ligase activity, we re-examined other moderately-conserved substrate-binding residues, particularly those in the S2’ pocket. PiAEP1 and PeAEP1 differ at position 172, which is a conserved Gly in 93% of legumains including PiAEP1. At this position, PiAEP1 has Gly, but PeAEP1 has Ser (corresponding to Ser169) (Supplementary Fig. S8). Mutation of Ser169 to Gly rescued part of the ligase activity to give a C/H ratio to 3.1 at pH 6.5, which represents a 6-fold increase relative to wild type (Supplementary Fig. S9). These results suggested that variations of the conserved Gly in or near the S2’ pocket could shift legumains towards hydrolysis. The replacement of Gly at this position could disturb the stability of enzyme-substrate interaction on the prime side. A recent mutagenesis study by Dall, et al. on human legumains also showed that V155G (homologous to butelase-1 Gly172) mutation could increase ligase activity.(Dall et al., 2021)

Overall, our functional study using the C/H ratios validated LADs of three legumain types. At their optimal pH, PALs have a C/H ratio of >20 and AEPs <0.25 — a 100-fold difference. The hybrid-legumain with C/H ratios <1 to 7.2 showed that the LAD2 at the prime side could also serve as a “gatekeeper” of ligase activity through Ala167 and residues in the S2’ pocket.

## DISCUSSION

An accepted mechanism for legumain catalysis is the formation of an Asx-thioester intermediate. However, this S-acyl intermediate is chemically unstable and prone to intramolecular cyclization with its side-chain amide or carboxylic acid to form, respectively, a succinimide or anhydride. In this regard, legumains have evolved a highly sophisticated substrate-binding pocket pattern, S2-S1-S1’-S2’, which not only stabilizes the thioester acyl intermediate to prevent intramolecular side-chain cyclization, but also facilitates intermolecular reactions to accept either an attacking water or amine nucleophile during a hydrolase or ligase reaction. This rationale forms a basis of our proposed molecular determinants in exploiting substrate-binding pockets for the legumain catalytic actions.

Four conserved glycine residues lie within or near the S2-S1-S1’-S2’ sites: Gly166 is invariant and Gly172 is also highly conserved in both AEPs and PALs. Our mutagenesis study suggested that Gly172 is a determinant for PALs to maintain a high C/H ratio. The remaining two Gly residues, Gly167 and Gly 237, are critical, but negative, determinants of ligase activity.

Previously, we suggested that amino acid residue 237 of LAD1 on the non-prime side functions as a “gate-keeper”. In this study, we show that Gly at position 237 occurs in 90% of legumains and is a negative ligase determinant. Substitution of LAD1-Gly237 with an aliphatic amino acid enhances ligase activity, and this increase correlates with the bulkiness of the substituted amino acid side chain. Thus, the gatekeeper role of LAD1 at position 237 is likely associated with thioester stabilization.

Unlike Gly237 of LAD1, Gly167 is less well defined because it depends on residue 168 and, to a certain extent, residue 172. Substitution of Gly167 with Ala facilitates ligase activity but substitution of other residues, such Ser and Tyr, promotes protease activity. Our mutagenesis study also suggested that prime-side LAD residues have greater influence on the directionality than those on the non-prime side. Thus, we propose the prime-side residue 167 could serve as a true “gate-keeper” for the incoming nucleophiles (Fig. 5). Brandstetter and co-workers proposed that retention of a leaving group or incoming nucleophile in prime-side pockets could exclude access by water and enhance ligation.(Zauner et al., 2018) Our results agree with this hypothesis by revealing the synergic effect of prime side residues at positions 167 and 172.

**Figure 5.**
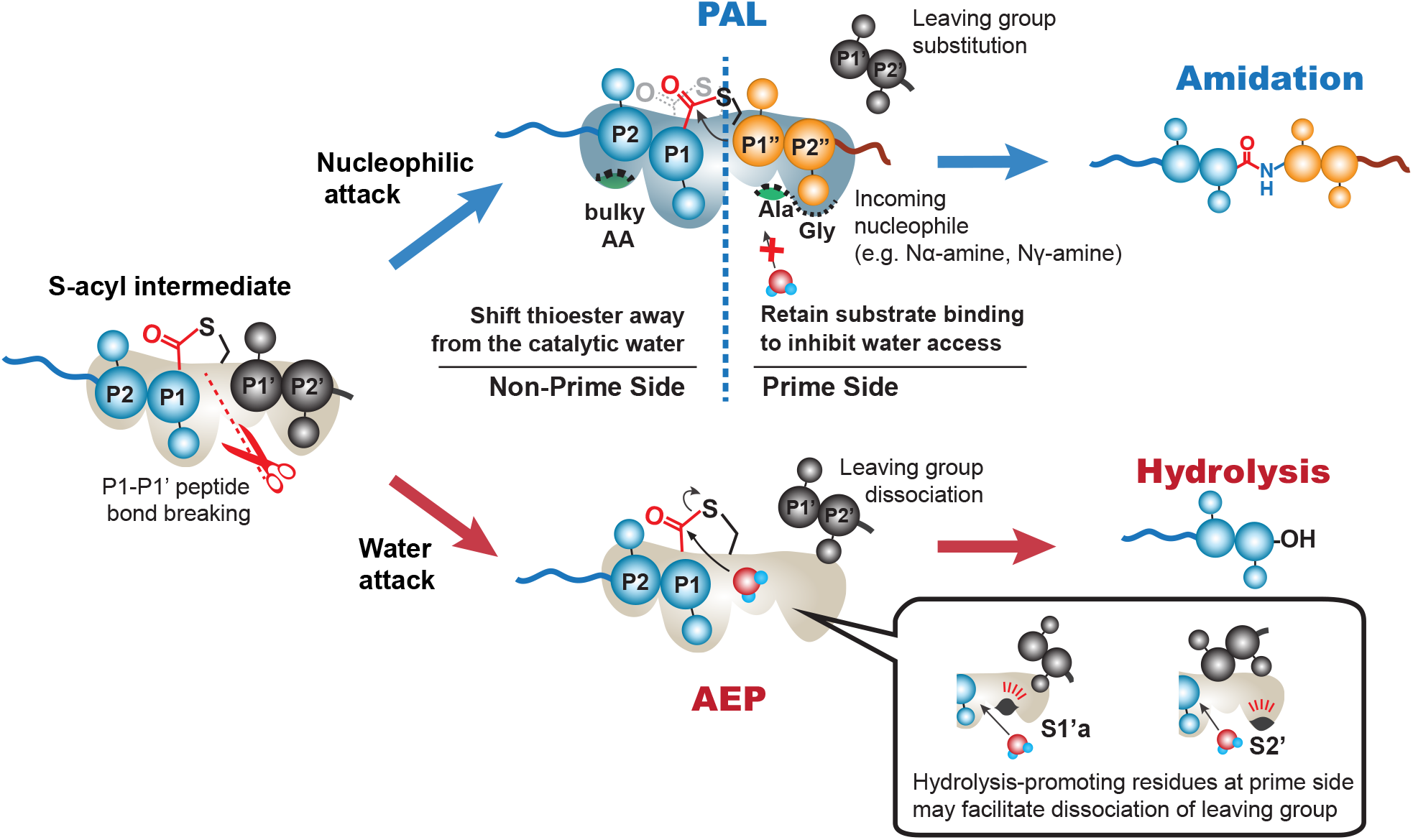
Proposed roles of LAD motifs based on functional and mutagenesis study. PAL-like LAD1 facilitates the nucleophilic attack by displacing the thioester intermediate away from catalytic water. PAL-like LAD2 favor peptide binding rather than water molecules. Similarly, conserved S2’-Gly172 also stabilizes enzyme-substrate interaction. AEP-like LAD2/S2’ mutations favor hydrolysis by facilitating a rapid substrate dissociation.

Expanding the repertoire of PALs will broaden their utility in biochemical and biotechnological applications. For example, the optimal and operational pH for PALs such as butelase-1, VyPAL2 and OAaEP1b is near neutral. This work and recent progress on PALs have extended their operational pH from pH 5.5 (e.g., BmAEP1-S161A) to pH 9 (e.g., McPAL1), and substrate P1 site recognition from Asn/Asp to unnatural amino acids such as Asn(OH) and Asn(Me) in addition to P2’ recognition signals. In turn, these advances allow various site-specific modifications as well as semi- and total-synthesis of proteins in tandem, orthogonally or under one-pot conditions.

In conclusion, plant legumains can catalyze hydrolysis and/or ligation of Asx-peptide bonds. In this study of 1,500 plant legumains, we exploited, analyzed and validated the usefulness of multiple sequence variants in the S2-S1-S1’-S2’ substrate-binding pockets as the molecular basis to distinguish hydrolase and ligase activity and for classification as PALs or AEPs. We observed that the enzymatic directionality of legumains could be “edited” by engineering key residues in substrate-binding sites. Similar principles could be applied to the engineering of other 123 subfamilies of thioester-forming Cys proteases, such as structurally similar caspases from the C14 subfamily, to evoke their ligase potentials.

## ASSOCIATED CONTENT

### Supporting Information

Supplementary files include materials and methods, additional experimental data (mass spectra, HPLC profiles, structural illustrations, gel images, summary tables), and supporting datasets.

### Author Contributions

JPT, XH and NYC designed the research. NYC and XH performed bioinformatic analysis, XH, NYC, LHT, and SH expressed and performed activity study of recombinant enzymes, XZ provided synthetic substrates, AS performed proteomic study. JPT, XH, and NYC wrote the manuscript. CFL and JL provided revision suggestions to the manuscript writing. #These authors contributed equally. All authors read and approved the final version of the manuscript.

### Funding Sources

This research was supported by the Academic Research Grant Tier 3 (MOE2016-T3-1-003) from the Singapore Ministry of Education and Nanyang Technological University.

## ACKNOWLEDGMENT

We thank Dr. Ka Ho Wong for his pioneer contribution to the bioinformatics analysis.

