## Supplemental Figures and Tables for "Substrate-binding Glycine Residues are Major Determinants for Hydrolase and Ligase Activity of Plant Legumains"

Hemu, et al.

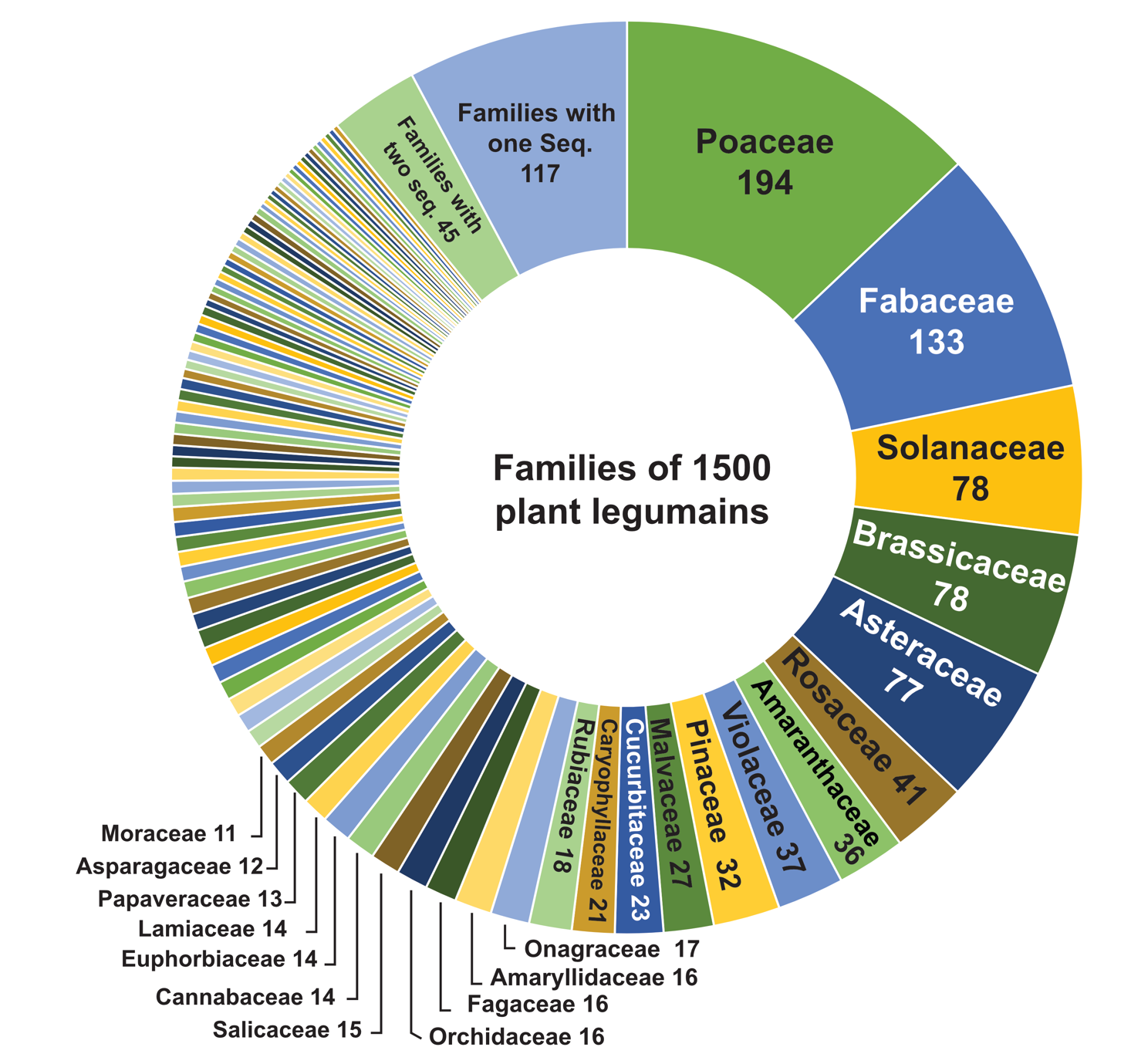

**Supplementary Figure 1.** Family distribution of 1500 plant legumain sequences in 249 plant families. See Dataset1 for the full list.

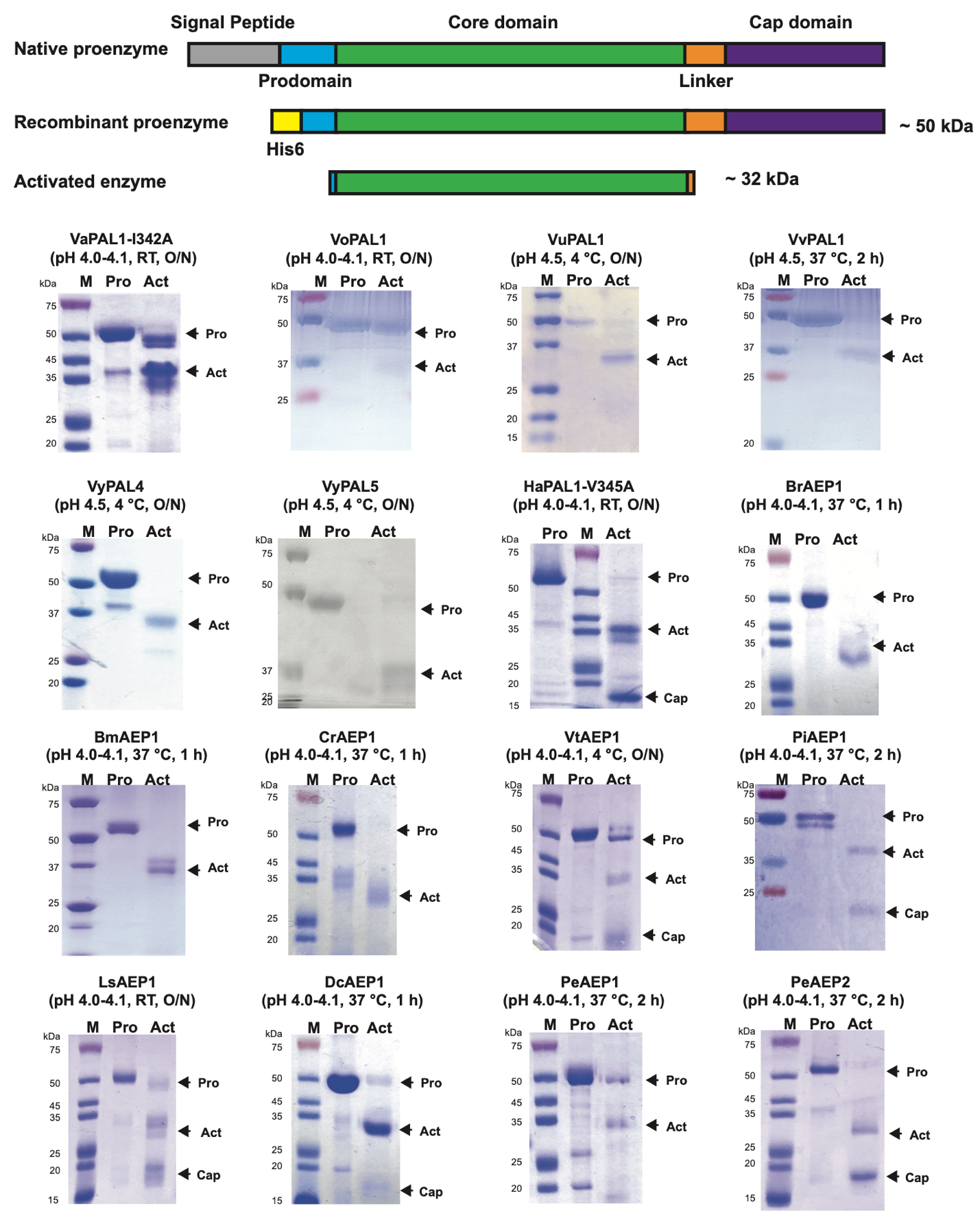

**Supplementary Figure 2.** Construct design and SDS-PAGE gels for 16 recombinant legumain proenzymes and activated forms. (**A**) Schematic representation of the His6-Pro-Core-Cap construct used in this study. (**B**) SDS-PAGE gels of purified proenzymes and activated enzymes. Conditions used are given in brackets. The bands of active enzymes appeared heavier than their actual masses due to the low isoelectric point (pI < 6) of activated enzymes.

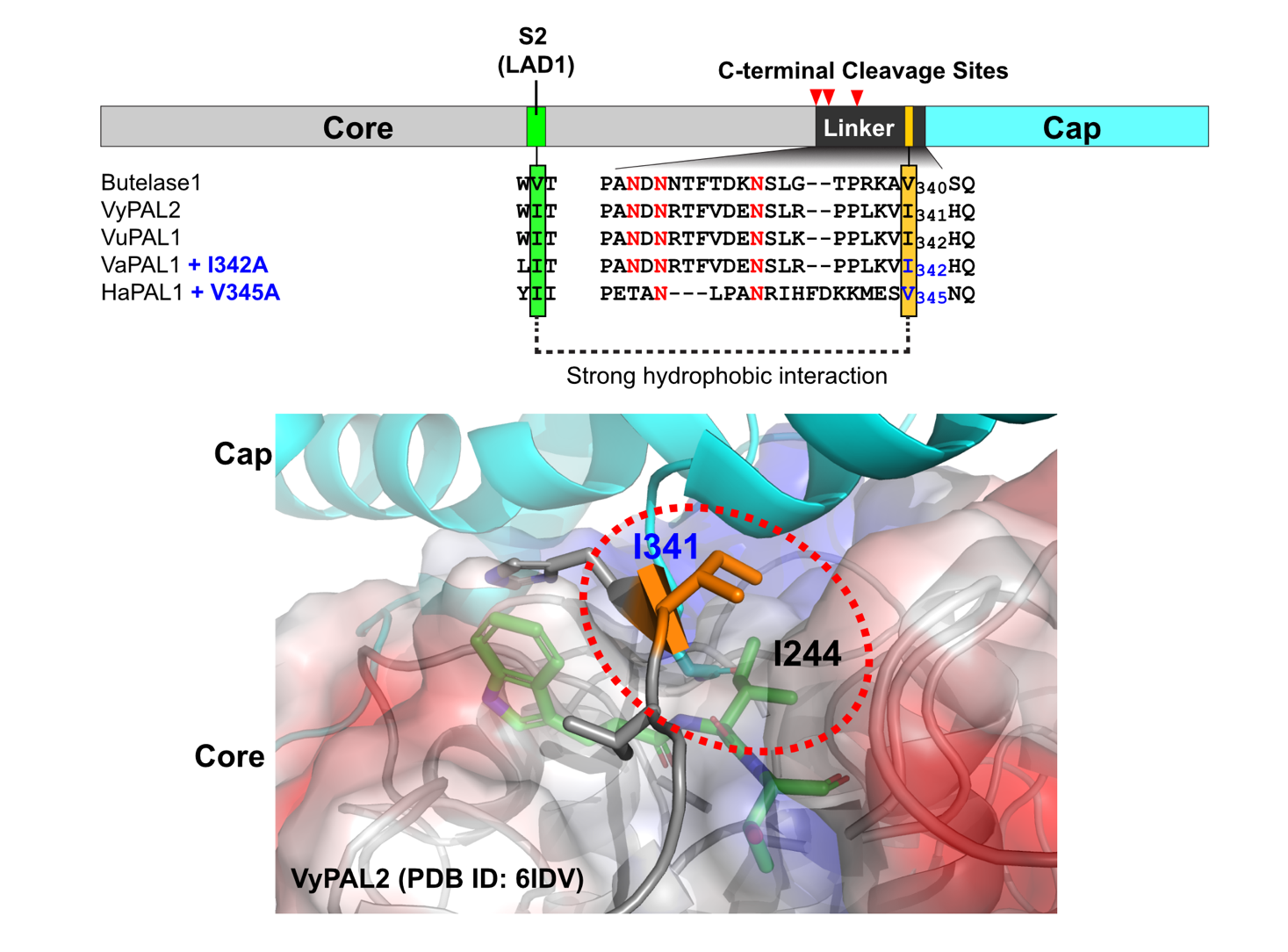

**Supplementary Figure 3.** Ala-substitution of the conserved hydrophobic residue in the linker region facilitate autolytic activation of VaPAL1 and HaPAL1 facilitated auto-activation likely due to the reduced hydrophobic interactions between linker and the hydrophobic S2 pocket that facilitated dissociation of cap domain from core domain.

| Core domain | Linker |
| --- | --- |
| **C-terminal cleavage site N333**  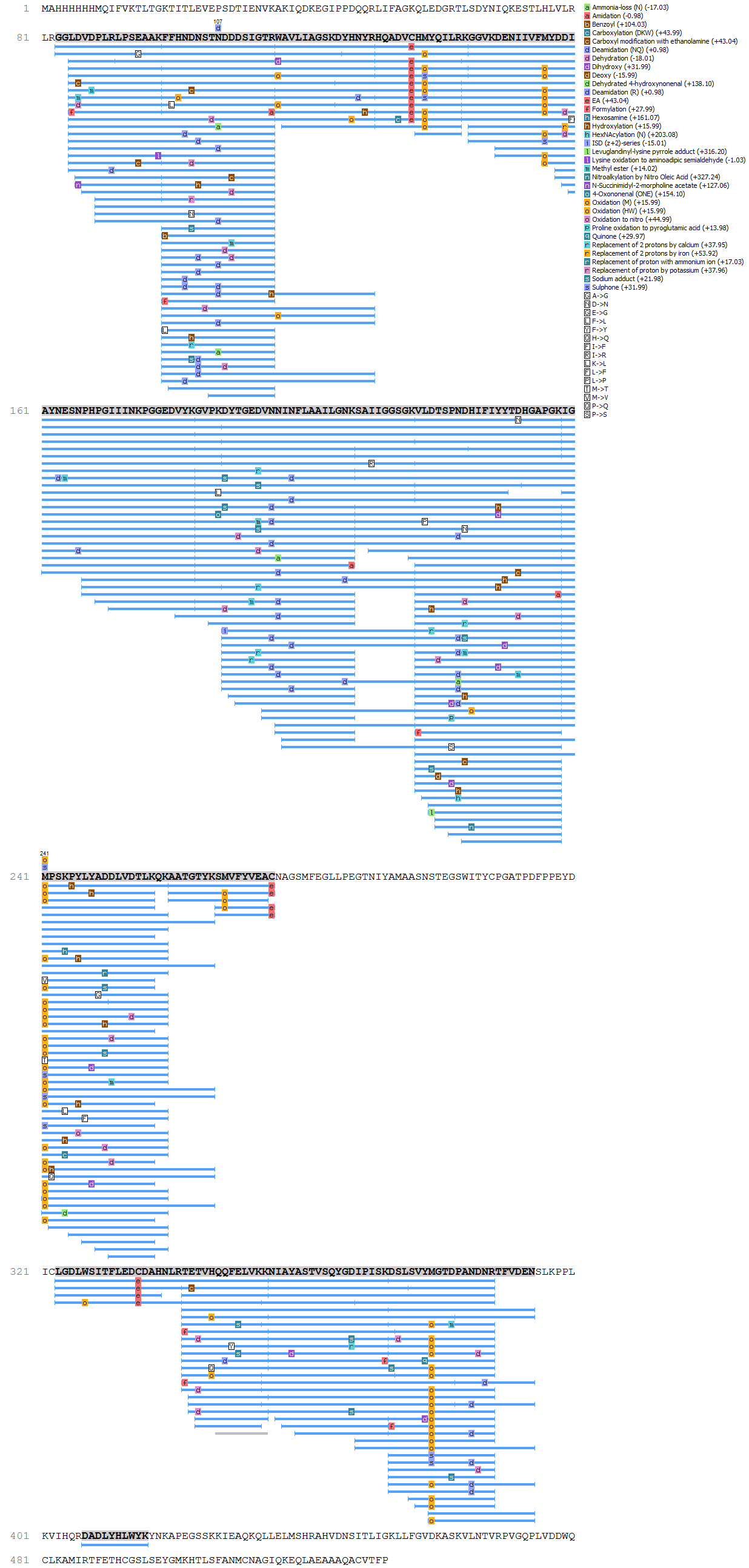 | |

**Supplementary Figure 4.** LC-MSMS sequencing of tryptic digested activated VuPAL1 shows the C-terminal cleavage site mainly at the conserved N333 in the linker region. Sequence after the cleavage site has been removed by autolytic hydrolysis.

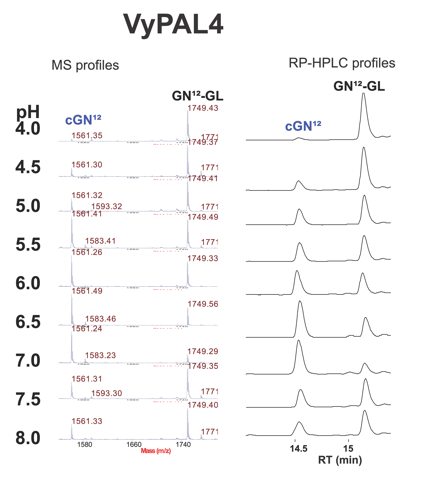

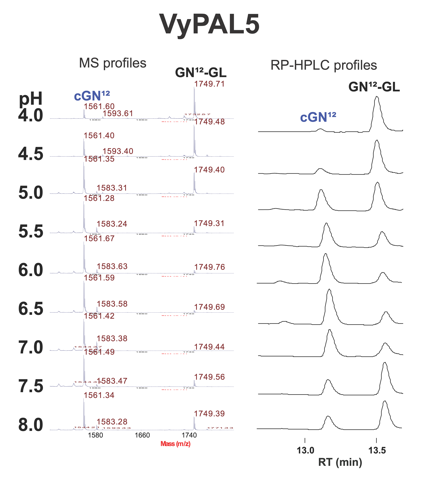

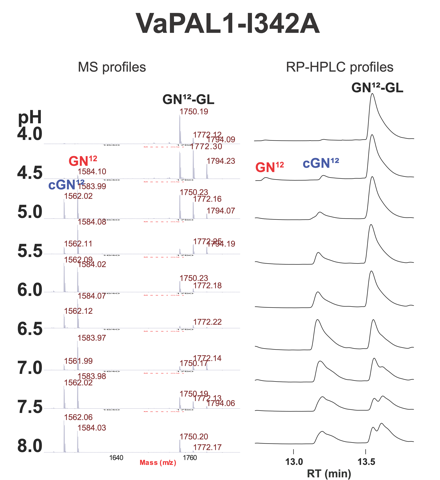

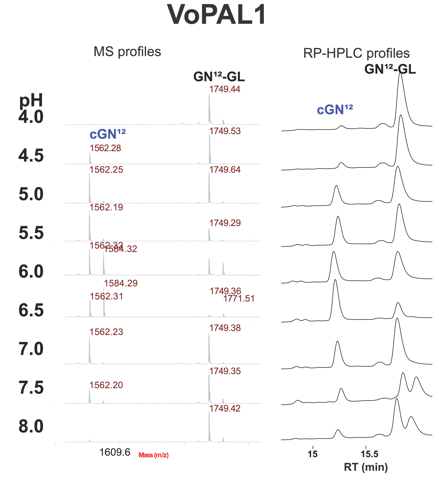

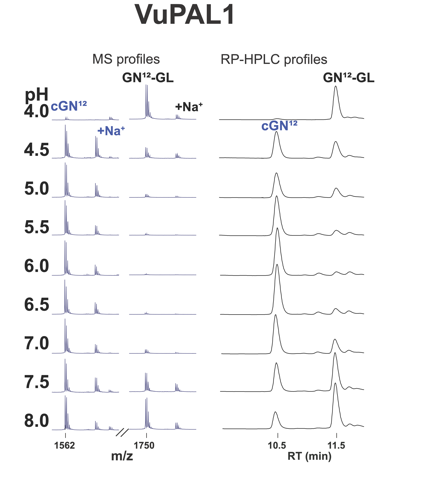

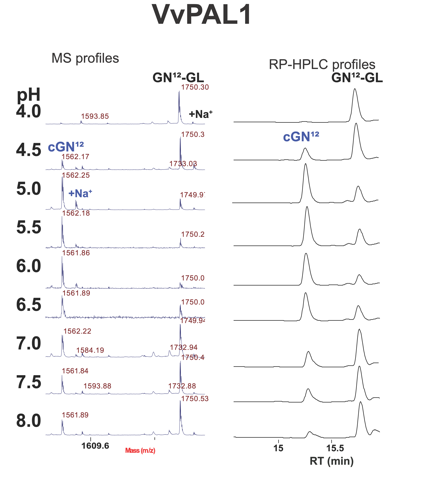

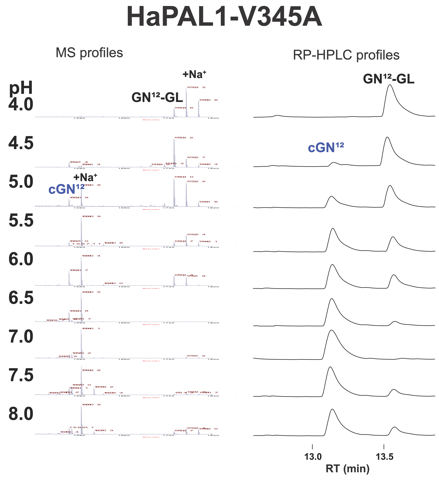

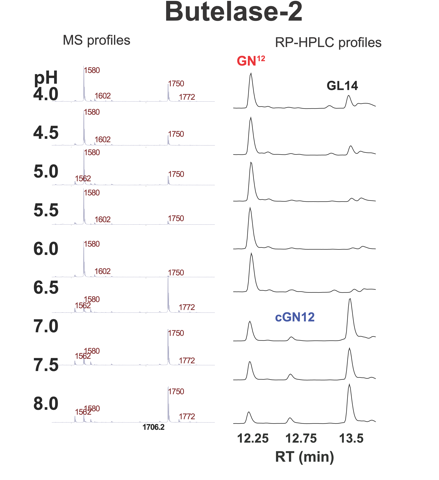

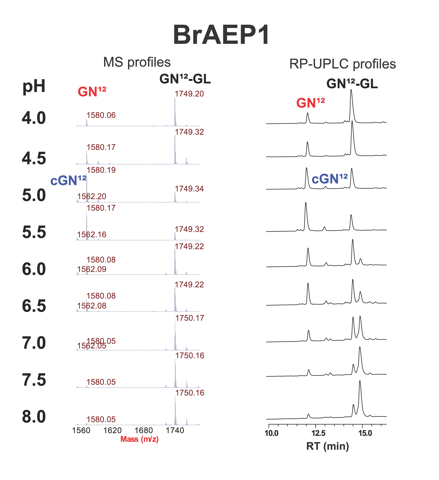

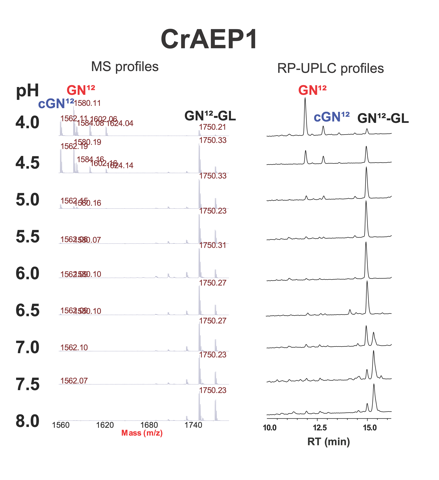

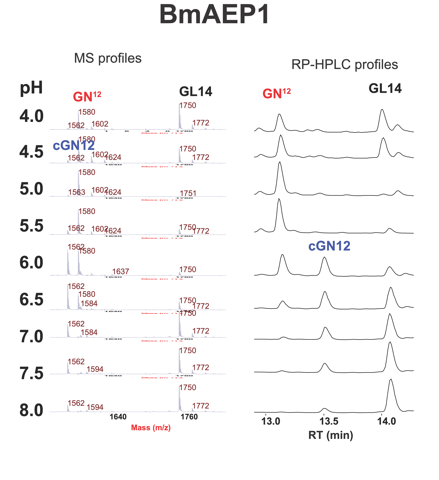

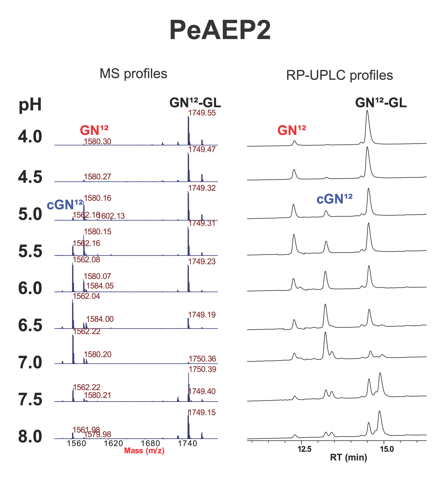

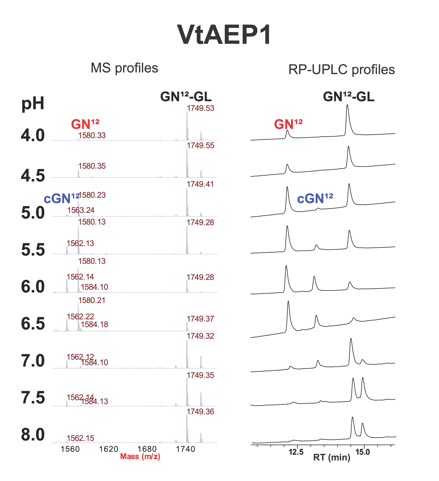

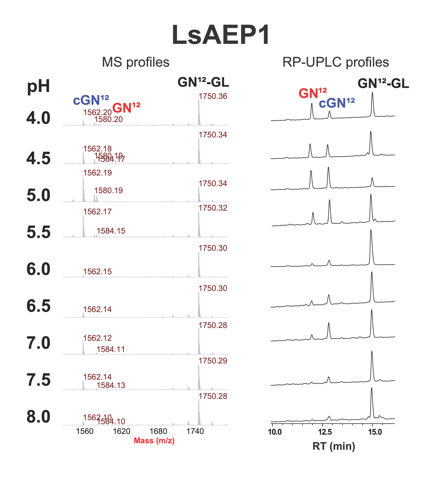

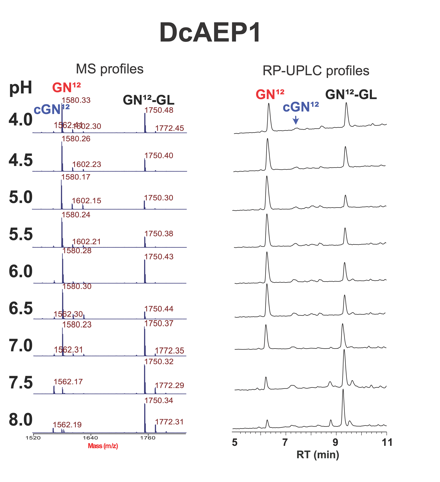

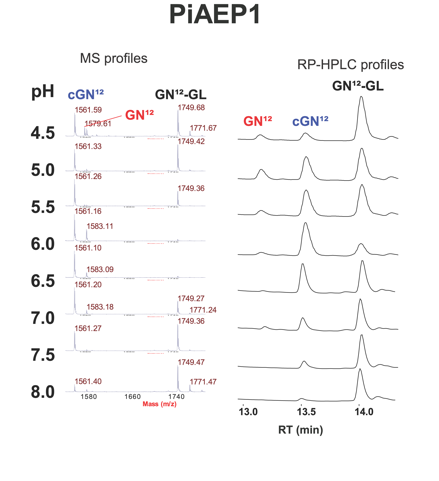

**Supplementary Figure 5.** MS and HPLC profiles of 16 recombinant legumains. One representative of the three repeated sets of reaction profiles is displayed for each enzyme. HPLC analysis were performed either on a 250x4.6 mm, 3.6 μm analytical column or a 150x2.1 mm, 1.7 μm UPLC column. We observed splitting peaks for the reaction mixtures performed at neutral to basic pHs, which could due to the change of ionization state of multiple Arg in the substrate sequence. We also observed that acidification of the reaction mixtures before injecting into HPLC/UPLC columns could resolve this problem.

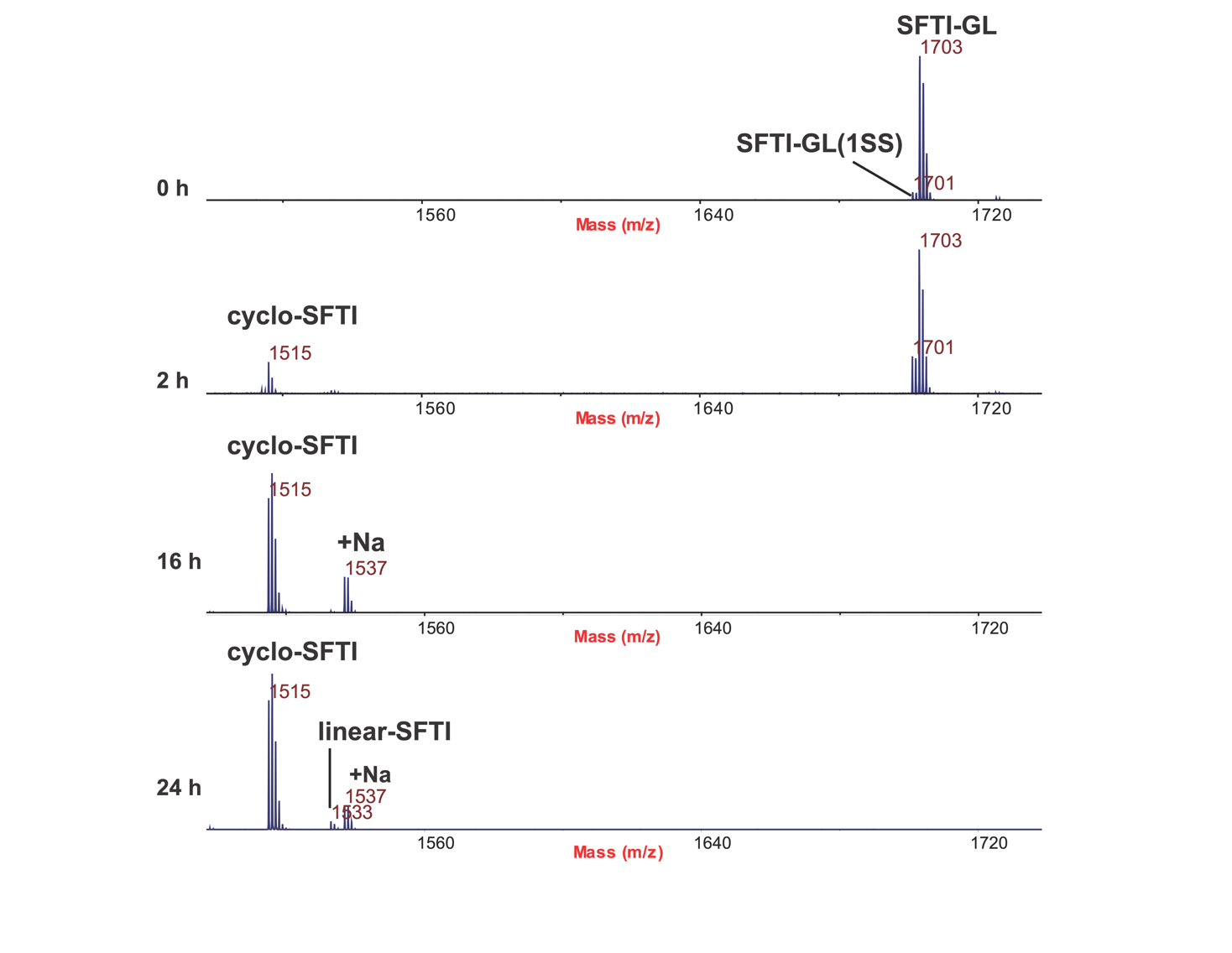

**Supplementary Figure 6.** MALDI-TOF mass spectrometry profiles of HaPAL1-mediated cyclization of SFTI-1. Reaction was performed at pH 6.0 at 37 °C with an enzyme : substrate molar ratio of 1:200. Hydrolytic product SFTI-1 (1533 Da) was observed after 24 h incubation.

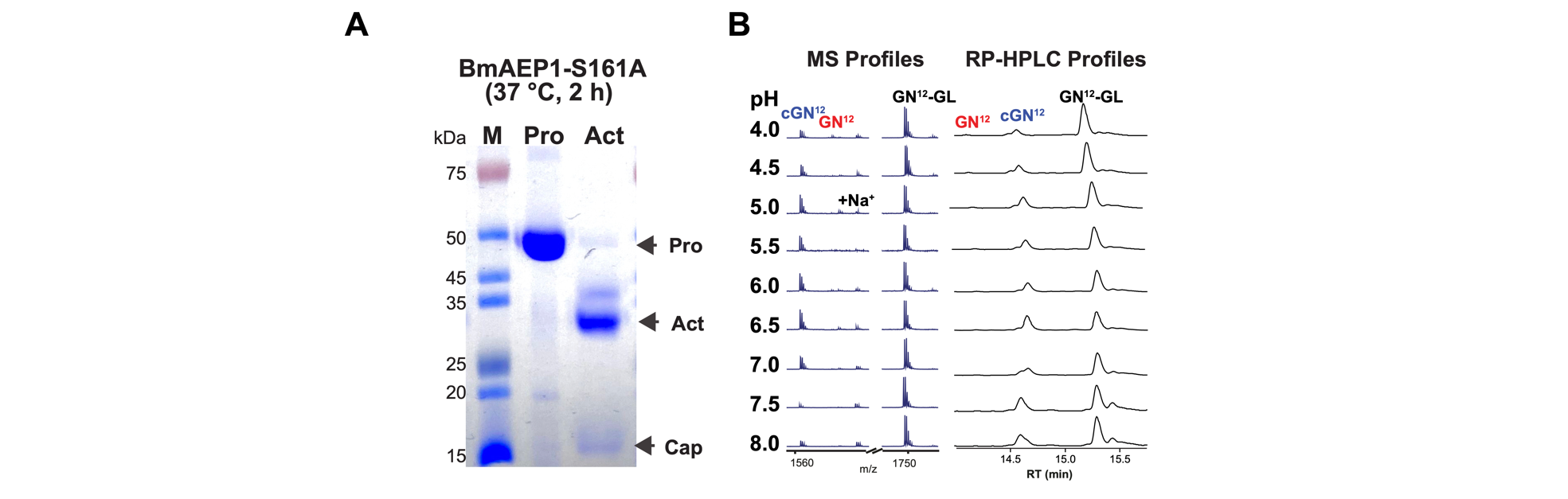

**Supplementary Figure 7.** SDS-PAGE gel, MALDI-TOF MS and HPLC profiles of BmAEP1-S161A.

- **Starting of the cDNA sequence for recombinant expression**

```
VyPAL4     ---------MKLLAAGVILVSLLALSGTVAVAVAGGLDVDP---LRLPSEAAKFFHNDNSTND--DDSIGTTWAVLIAGSKGYHNYRHQADVCHMYQILRKGGVKDENIIVFMYDDIAYNESNPFPGIIINKPGGENVYKGVPKDYTGEDINNVNFLAAILGNKSAIIG-GSGKVL 161
VyPAL5     ---------MKLLAAGVILVSLLALSGTVAVAVAGGLDVDP---LRLPSEAAKFFHNDNSTND--DDSIGTTWAVLIAGSKGYHNYRHQADVCHMYQILRKGGVKDENIIVFMYDDIAYNESNPFPGIIINKPGGENVYKGVPKDYTGEDINNVNFLAAILGNKSAIIGGSGKVL 161
VuPAL1     ---------MKLLAAGVILVSLLALSG----TVAGGLDVDP---LRLPSEAAKFFHNDNSTND--DDSIGTRWAVLIAGSKDYHNYRHQADVCHMYQILRKGGVKDENIIVFMYDDIAYNESNPHPGIIINKPGGEDVYKGVPKDYTGEDVNNINFLAAILGNKSAIIGGSGKVL 157
VaPAL1     ---------MKLFAAGVILFSLLALSG----TIAGGLDVYS---LRLPSEAAKFFHNDNSTND--DDSIGTRWAILIAGSKGYNNYRHQADVCHMYQILRKGGVKDENIIVFMYDDIAYNESNPHPGILINKPGGEDVYKGVPKDYTGEDINNVNFLAAILGNKSAIIGGSGKVL 157
VoPAL1     ---------MKLLAAGVILFFLLALSG----TVAGGLDVDP---LRLPSEAAKFFHNDNSTND--DDSVGTRWAVLIAGSKGYQNYRHQADVCHMYQILRKGGVKDENIIVFMYDDIAYNESNPHPGILINKPGGEDVYKGVPKDYTGEDINNINFLAAILGNKSAIIGGSGKVL 157
VvPAL1     ---------MKLFAAGVILFFLLTLSG----TIAGGLDVDP---LRLPSEAAKFFHNDNSTND--DDSIGTRWAVLIAGSKGYHNYRHQADVCHMYQILRKGGVKDENIIVFMYDDIAYNESNPHPGILINKPGGEDVYKGVPKDYTGEDINNINFLAAILGNKSAIIGGSGKVL 157
HaPAL1     ----MACFSYRLICLLLVLMMVMALPNGAAAARR--GSDYWDPFIRSPVDLE---DDELGN--------GTRWALLVAGSKGYQSYRHQANVCHAYQILKRGGLKDENIVVFMYDDIATCDENPRPGTIIHHPEGGDVYAGVPKDYTGDAVTADNFFAVILGDKSSVKGGSGKVI 158
Butelase-2 MAVDHCFLKKKTCYYGFVLWSWMLMMSLHSKAARLNPQKEWDSVIRLPTE-----PVDADT-----DEVGTRWAVLVAGSNGYENYRHQADVCHAYQLLIKGGLKEENIVVFMYDDIAWHELNPRPGVIINNPRGEDVYAGVPKDYTGEDVTAENLFAVILGDRSKVKGGSGKVI 166
BrAEP1     --------MTPVPV----AVIFLSLIAL-----SAGRQNPDDDVIKLPSQASRFFRPNNDD---ESSPSGTRWAVLVAGSSGYWNYRHQADVCHAYQLLRKGGLKEENIVVFMYDDIADNEENPRKGIIINSPHGSDVYEGVPKDYTGDDVTVENLFAVILGDKGAVKGGSGKVV 155
CrAEP1     -----MAVYSALFTVTLALLLVVSLPLKASGSRKASPG-HWDPFIQSPSDE-----EDNR---------GTRWAILVAGSNGFDNYRHQSDVCHAYQILKNGGLKDENIVVFMYDDIANNKLNPRPGIIINHPDGEDVYEGVPKDYTGEHVTPENLFAVLLGDESSLKGGSGKVV 155
BmAEP1     --------MATCYATSTKFVLLIALLLFS------------------DIIAKRESVDGAS-----TDQPGKRWAILVAGSSGYENYRHQADVCHAYQILRKGGLPDENIIVFMYDDIAFNPSNPRPGVVINKPDGVDVYQGVPKDYTGEHVNSINFYAVILGNRSALTGGSGKVV 144
PiAEP1     ---------MARYAG-VAALLLLLALSA-----VAVSGSR-DDFLKLPSEVADFFRHKASNDGVDFDSVGTRWAVLIAGSKDYGNYRHQADVCHAYQILKRGGLKDENIIVFMYDDIAYSEDNPRPGVIINGPNADDVYEGVPKDYTGDEVNAKNFLAAILGDRSAITGGSGKVV 159
LsAEP1     -----------MSAFHFALLLLTLLTVKLSEATR-------SSLFASTKDVN-----------------STKWAVLVAGSSGYYNYRHQADVCHAYQILKKGGLKDENIIVFMYDDIASNIMNPRPGVIINSPNGSDVYAGVPKDYTGGYVTVANFYAVLLGNASGVTGGSGKVV 140
DcAEP1     ----------------------------------------------------------------DDDTVGTKWAVLIAGSDGYWNYRHQADICHAYQLLKKGGLKDENIVVFMYDDIAYNEENPRQGVIINSPYGSDVYAGVPKDYTGKDVNANNFIAALLGDKAALTGGSGKVV 111
PeAEP1     ----------MVQKYGGTTLFLVALFVL----------------AVCTAEARSLLHEISNANH--DNSIGTKWAVLVAGSNYWFNYRHQADVCHAYQLLKQGGLKDENIIVFMYDDIAYNKENPRPGVIINSPHGENVYEGVTKDYTGEHCNADNFFAVILGNKTALTGGSGKVV 147
PeAEP2     -----------MISHVAGILILVGFSI--------LGAGEGRDVLKLPSEASRFFKKGE-----DDDSVGTRWAILLAGSNDYWNYRHQADVCHAYQLLRKGGLKDENIVVLMYDDIAYNEENPRKGVIINNPAGEDVYKGVPKDYTGDDVNVDNFLAVLLGNKTAITGGSGKVV 151
VtAEP1     -------------MAALALLFLLALSG----FVSGGRDITGDESLPLP-----SFQGNRNSDDDVDSSAGTKWAVLIAGSKGYQNYRHQADVCHAYQILRRGGVKDENIIVFMYDDIAYHIRNPYPGTITNSPDRKDVYKGVPKDYTGEDVNVQNFLAVILGNKTALTGGSGKVL 153

                         H LAD2 S2’-Gly                             C                             LAD1	poly-Proline			       MLA
VyPAL4     DTSPNDHIFIYYADHGAPGKIGMPSK-PYLYADDLVDTLKQKAAAGTYKSMVFYVEACNAGSMFEGLLPEGMNIYAMTASNSTEGSLIAYCAGV-----TPGVP--LEIVTCLGDLWSITFLEDCDAHNLRTETVHQQFELVKKK------IAYASTVSQYGDIPISKDSLSVYM 322
VyPAL5     DTSPNDHIFIYYADHGAPGKIGMPSK-PYLYADDLVDTLKQKAATGTYKSMVFYVEACNAGSMFEGLLPEGTNIYAMAASNSTEGSWITYCPG------TPDFP--PEFDVCLGDLWSITFLEDCDAHNLRTETVHQQFELVKKK------IAYASTVSQYGDIPISKDSLSVYM 321
VuPAL1     DTSPNDHIFIYYTDHGAPGKIGMPSK-PYLYADDLVDTLKQKAATGTYKSMVFYVEACNAGSMFEGLLPEGTNIYAMAASNSTEGSWITYCPGA-----TPDFP--PEYDICLGDLWSITFLEDCDAHNLRTETVHQQFELVKKN------IAYASTVSQYGDIPISKDSLSVYM 318
VaPAL1     DTSPNDHIFIYYADHGAPGKIGMPSK-PYLYADDLVDTLKQKAAAGTYKSMVFYVEACNAGSMFEGLLPEGMNIYAMTASNSTEGSLITYCAGV-----TPGVP--LEIVTCLGDLWSITFLEDCDAHNLRTETVHQQFELVKKR------IAYASTVSQYGDIPISKDSLSVYM 318
VoPAL1     DTSPDDHIFIYYTDHGAPGKIGMPSK-PYLYADDLVDTLKQKAATGTYKSMVFYVEACNAGSMFEGLLPEGVNIYAMAASNSTEGSWVTYCPG------TPDFP--PEFDVCLGDLWSITFLEDCDAHNLRTETVHQQFELIKKK------IAYASTVSQYGDIPISNGSLSVYM 317
VvPAL1     DTSPDDHIFIYYTDHGAPGKIGMPSK-PYLYADDLVDTLKQKAATGTYKSMVFYVEACNAGSMFEGLLPEGTNIYAMAASNSTEGSWITYCPG------TPDFP--PEFDVCLGDLWSITFLEDCDAHNLRTETVHQQFELIKKK------IAYASTVSQYGDIPISNDSLSIYM 317
HaPAL1     DSKPDDRIFLYYTDHGAAGLLGMPEK-PYVVANDFVEVLKKKHAMGTYKEMVIYLEACESGSIFEGLLPEDLNIYAITSTKPEEPSYIIYCP-------DMNPPPPPEYTTCLGDTFSVAWMEDSETHNLKKESLAQQINKVKERTSMFGTYANGSHVMEYGTKVIKPEKVYLYQ 325
Butelase-2 NSKPEDRIFIFYSDHGGPGVLGMPNEQ-ILYAMDFIDVLKKKHASGGYREMVIYVEACESGSLFEGIMPKDLNVFVTTASNAQENSWGTYCP-------GTEPSPPPEYTTCLGDLYSVAWMEDSESHNLRRETVNQQYRSVKERTSNFKDYAMGSHVMQYGDTNITAEKLYLFQ 332
BrAEP1     DSGPNDHIFIFYSDHGGPGVLGMPTS-PYLYADDLNDVLKKKHASGTYKSMVFYLEACESGSIFEGLLEEGLNIYATTASNAVESSWGTYCPG------EEPSPP-PEYETCLGDLYSVAWMEDSGMHNLQTETLRQQYELVKRRTAGGA-SAYGSHVMQYGDVGLNKDKLDLYM 321
CrAEP1     KSSHNDRIFIYYTDHGGPGVLGMPTRSGFLYAKDFIEVLKKKHASKTYKEMVIHVEACESGSFFEGLMPEDLNIYVQTASNADESSYATYCPG------SDPHS-PPDDFPCLGDLYSVAWMEDSESHNLKKETIVQQYKKVKDRTSNHKSYEGGSHVMQYGNKSINIEKLYLYQ 323
BmAEP1     DSDLHDHIFIYYTDHGSAGLLGMPEG-DYVYAKDLMEVLKQKHEAKSYKSMVIYVEACESGSMLEGLLPENIKIYATTASNATENSWATYCPGQ-----FPSPP-TDYDTCLGDLYSIAWMEDSDKHDLSKETLIQQYDAVRRRTLVDK-FGYGSHVMLYGNKSIGNN-SLDTYI 310
PiAEP1     DSGPNDHIFIYYTDHGAPGVIGMPKR-PYLYAHELIETLKKKHASGTYKSLVFYLESCEAGSMFEGLLPEDLNIYAFTASNAEESSWAAYCPGQ-----AHGSPP-PEYDICVADLYSVAWMEDSEVHNLRTETLRQQYGVVYKRNLD----ISGSHAMRYGDLNLSAEDLFLYM 323
LsAEP1     ASKPGDKIFVFYSDHGAPGILGMPTV-PHLYANDFIEVLKMKNASGTYDKMVIYIESCESGSIFEGLLPEDMNIYVTTASNANESSWGTYCP-------DMTPPPPPEFHTCLGDLYSISWMENSDLEDLTIETLEQQYSKVKIRTLNNNTEE-GSHVMQYGTQHISKETVSTYQ 306
DcAEP1     DSGPNDHIFVFYSDHGGAGVVGMPSY-PYLYADDLIDALKKKHALGSYKSLVFYLEACESGSMFEGILPDGLNIYATTASNAWESSWGTYCPG------EDAIP-PEYDTCLGDLYSISWMEDSEKRNLRAETLKQQYELVKKRTGSN--SFYGSNVMQYGDLGLNKD-VLYTYM 275
PeAEP1     NSGPNDHIFIYYADHGAPGMISMPN--DMIFADDLIKVLTKKNLDEAYRKLVFYLEACESGSMFDGLLPKGLNIYVTTASNPYESSWATYCSADGDEGCIGECPPKDFKDVCLGDLYSVSWLEDSDLHNRQVETLEQQYQVVRKRTLNNN-TQEGSHVMQYGDLHLSKDALFGYM 319
PeAEP2     DSGPNDHIFIFYTDHGGPGVLGMPTK-PYLYASDLIGALKKKHASGTYKSLVLYVEACEAGSIFEGLLPEGLNVYATTASDAVEGSWVTYCPG------QNPSPP-PEYTTCLGDLYSVSWMEDSEKHNLQTESLRQQYHLVKRK------IAYASHVMQYGYLKLSMDSLSMYM 312
VtAEP1     NNAPNDHIFIYYTDHGYPGVLGMPTE-PYLYANDLIDTLKKKHASGTYEAMVFYVEACESASMFEGLLPDGLNIYVSTAAKAGEGSWVTYCPTQ-----HPAVP--AEYGTCVGDLYSVTWMEDCDVYNLRTQTLHQQYEMVKKK------IAYASSVTQFGDLPISKDSLYKYM 314

VyPAL4     GTDPANDNRTFVD-----ENSLRPPL--KVIHQRDAYLYHLWYKYQNTPEGSSKKIEAQKQLLEMMSHRAHVDNS ITLIGKLLFGMDKASKMLNSVRPAGQPLVDDWQCLKTMIRTFERHCGSLSEYGMKHTLSFANMCNAGIRKEQLAEAAAQACVTFPSNSYSSLAEGFSA- 488
VyPAL5     GTDPANDNRTFVD-----ENSLRPPL--KVIHQRDAYLYHLWYKYQNTPEGSSKKIEAQKQLLEMMSHRAHVDNS ITLIGKLLFGMDKASKMLNSVRPAGQPLVDDWQCLKTMIRTFERHCGSLSEYGMKHTLSFANMCNAGIRKEQLAEAAAQACVTFPSNSYSSLAEGFSA- 487
VuPAL1     GTDPANDNRTFVD-----ENSLKPPL--KVIHQRDADLYHLWYKYNKAPEGSSKKIEAQKQLLELMSHRAHVDNS ITLIGKLLFGVDKASKVLNTVRPVGQPLVDDWQCLKAMIRTFETHCGSLSEYGMKHTLSFANMCNAGIQKEQLAEAAAQACVTFPSNSYSSLAEGFSA- 484
VaPAL1     GTDPANDNRTFVD-----ENSLRPPL--KVIHQRDADLYHLWYKYQNIPEGSSKKIEAQKQLLELMTHRAHVDNS ITLIGKLLFGMDNPSKMLNSVRPAGQPLVDDWQCLKAMIRTFETHCGSLSEYGMKHTLSFANMCNAGIRKEQLAEAAAQACVTFPSNSYSSLAEGFSA- 484
VoPAL1     GTDPANDNRTFVD-----ENSLRPPL--KVIHQRDADLYHLWYKYQNTPEGSSKKIEAQKQLLEMMSHRAHVDNS ITLIGKLLFGMDKASKMLNSVRPAGQPLVDDWQCLKAMIRTFETHCGSLSEYGMKHTLSFANMCNAGIRKEQLAEAAAQACVTFPSNSYSSLAEGFSA- 483
VvPAL1     GTDPANDNRTFVD-----ENSLRPPL--KVIHQRDADLYHLWYKYQNTPEGSSKKIEAQKQLLEMMSHRAHVDNS ITLIGKLLFGMDKASKMLNSVRPAGQPLVDDWQCLKAMIRTFETHCGSLSEYGMKHTLSFANMCNAGIQKEQLAEAAAQACVTFPSNSYSSLAEGFSA- 483
VvPAL2     GTDPANDNRTFVD-----ENSLRPPL--KVIHQRDADLYHLWYKYQNTPEGSSKKIEAQKQLLEMMSHRAHVDNS ITLIGKLLFGMDKASKMLNSVRPAGQPLVDDWQCLKAMIRTFETHCGSLSEYGMKHTLSFANMCNAGIQKEQLAEAAAQACVTFPSNSYSSLAEGFSA- 483
HaPAL1     GYNPETAN--LPANRI------HFDKKMESVNQRDGDLIYLWQKYKRSS--VSNRAEALKQMTETLRYMAHLDSS VDMIGVLLFGPQNGGSILRSSRGRGLPLVDDWDCLKSMTRLFEKHCGLLTEYGMKHMRAFANICNNLVEETEVEEAIIATCSGKNIG-PYASLGAYSV- 487
Butelase-2 GFDPATVN--LPPHNG------RIEAKMEVVHQRDAELLFMWQMYQRSNHLLGKKTHILKQIAETVKHRNHLDGS VELIGVLLYGPGKGSPVLQSVRDPGLPLVDNWACLKSMVRVFESHCGSLTQYGMKHMRAFANICNSGVSESSMEEACMVACGGHDAGHL---------- 488
BrAEP1     GTNPANDNFTFVDA-----NSLTPPSG--VTNQRDADLVHFWDKYRKAPEGSTRKTEAQKQVLEAMSHRLHVDNS VKLIGKLLFGISEGSEVLNKVRPAGQPLADDWTCLKNMVRAFERHCGSLSQYGIKHMRSFANICNAGIQMRQMEEAASQACTSIPSGPWSSLHRGFSA- 487
CrAEP1     GFDPTTEN--LPADNR------LPDAPMGVVEQRSADLFFLWKKYKKMENGKVEKAELLKQLTEKMLHRTHVDGS MQIIEAFLFGPGRTPSVFSFVREHGLPIVDDWGCLKSMVRIFETHCGPLNQYGMKYMRAFANICNHKVTQASMEEASIAACSGRKQETWNPSYLHFSA- 488
BmAEP1     GANPDNYNYTSSVQSTDTIAPPSKLLYSNAVSQRDASLIHYWHKFQKAPFGSREKTEARKQLEDEILNRRHVDSS IYHIAKLLFGQAKSSEVLNNVRQQGQSLVDDWGCFKKFVKTYEKHCRRLSRYGMKYTRALANICNAGITINQMDQACLETCLVKT-------------- 470
PiAEP1     GSNPANDNSTFVDD-----NALRPFS--KAVNQRDAELVHFWEKFKRAPEGSLRKLKAQKEVFEAMSHRMHVDDS IKLIGKLLFGIEKGPEILKAVRPAGQPLVDDWDCLKALIRTFETHCGSLSQYGMKHTRSIANICNAGIKTEQMAEAAAQACVSVP-------------- 476
LsAEP1     G--SSTWN---TTTNS-----IISLGSMGVVDQRVADLYSMWQTYEKSTGEPQEKIELLKNIKEITTHRAHLDSS VETIKGKLGDQDYG-----SVRPEGSVLVDDWECLKSMIRTFETHCGSLTQYGLKHSRTFANMCNNGVTKEAMDEASKATCSSFNMGQWNPATVGYSA- 464
DcAEP1     GTNPANDNITNVDN-----NSLHPTSS-VAVNQRDADLIHFWSKLRKAPEGSARKVEAQKKLLEAISHRVHLDNS VQIIGKLLFGTEKSSEMLQTIRPAGQALVDDWKCLKSQVRTFETHCGSLSQYGMKHMRSFANICNAGVTTEQMAEAAAQACPSFPSNPWSSLHKGFSA- 442
PeAEP1     GSNSSTKN-----------HESRLSS--KMINQRDVHLWYLRSKFQSAPEGSARKIEASRQLNEAIAQRKHVDDS VRHIGELLFGVEKGQEVLKTIRPAGESLVDDWDCLKSFVKTFEEHCGKLTPYGRKHVRGFANLCNAGIQREQMDAAAKQACAL---------------- 464
PeAEP2     GTDPANDNYTFVDD-----NSLGASS--EAVNQRDADLLHFSDKFLKAPEGSARKVEAQKQFAEAMSHRMHLDNS MALVGKLLFGIKKGPEVLKRVRSDGQLLVDDWACLKSFVRTFETHCGSLSQYGMKHMRSIANICNAGIEVEQMVEASSHACPSIPSNTWSSLHRGFSA- 478
VtAEP1     GTDPSNDKHQYVDQ----ENSLKAHV--DAVHQREADLYHFWDKYQKASEGSRNKIDARKQLVEVMLHRMHVDDS IES----------------TIRPAGQPLVYDWDCLKTMVRTYETHCGSLSEYGMKYTRFLANICNSGVEKEKMAEAAAQVCVNFPSNPWSSLVKGFSA- 465
```

**Supplementary Figure 8.** Sequence alignment of 18 legumains. Catalytic dyad His and Cys are shaded in yellow. Residues corresponding to position 167, 172 and 237 are highlighted in red.

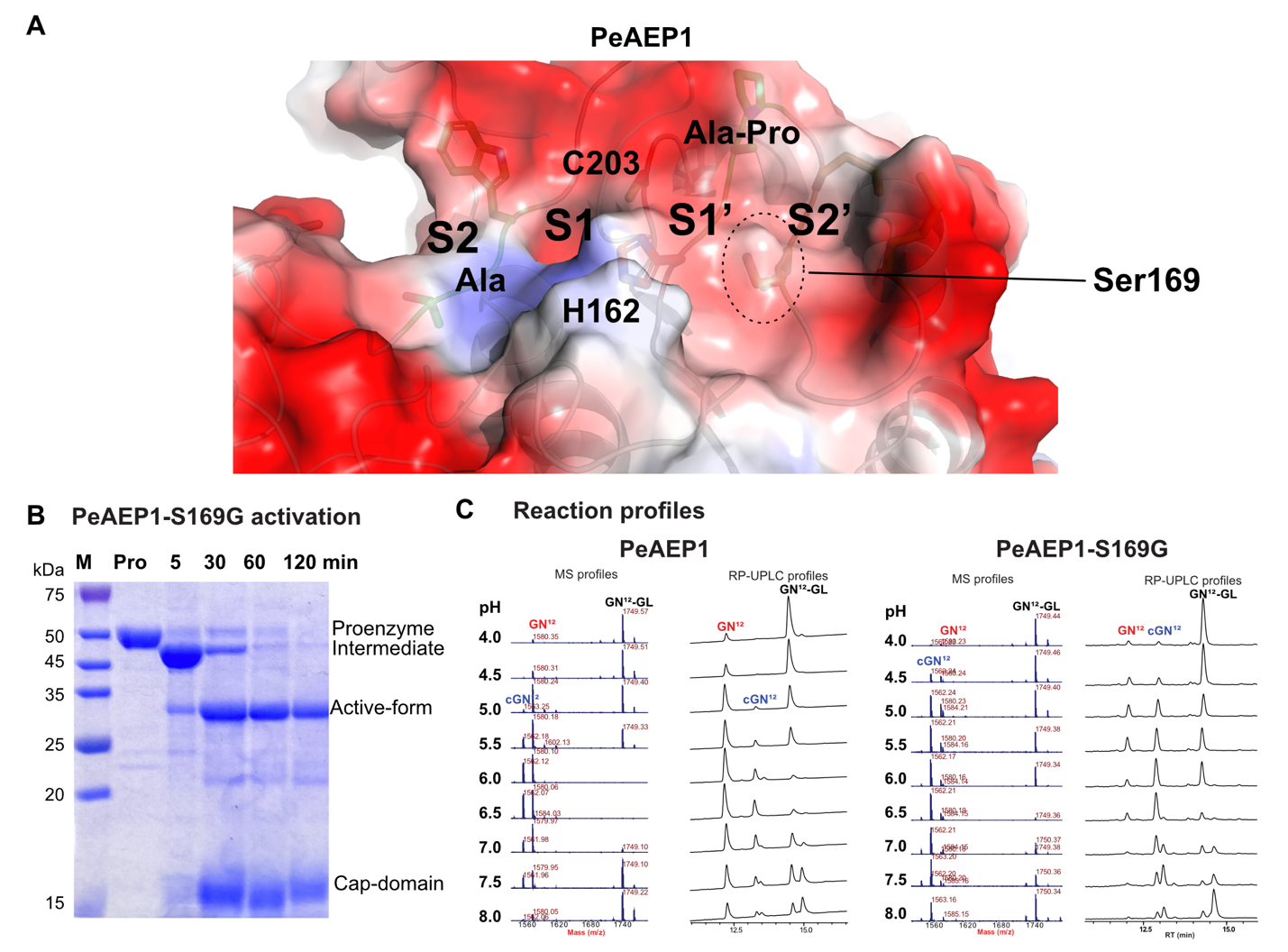

**Supplementary Figure 9.** Experimental data of PeAEP1 and PeAEP1-S169G. (A) Mapping of the catalytic dyad H162/C203 and LAD residues on S2-S1-S1’-S2’ pockets on the modelled PeAEP1 3D structure. The hump at the bottom of S2’ pocket posed by Ser169 is circled with a dashed line. (B) SDS-PAGE result for activation of PeAEP1-S169G. Activation conditions performed for 5, 30, 60 or 120 min were at 37 °C. Overnight (O/N) activation was performed at room temperature (25 °C) for 16 h. (C) Examples of MALDI-TOF MS and HPLC profiles of PeAEP1 and PeAEP1-S169G, respectively.

**Table S1. Amino acid composition of LADs in 1495 legumains.**

| Location | Composition | No. of sequences | Ratio (%) |
| --- | --- | --- | --- |
| Gly237 | **G** | 1346 | 89.7 |
|  | **A** | 121 | 8.1 |
|  | **V** | 15 | 1.0 |
|  | **I** | 12 | 0.8 |
|  | **C** | 3 | 0.2 |
|  | **P** | 2 | 0.1 |
|  | **S** | 1 | <0.1 |
| Gly167-Pro168 | **GP** | 1259 | 83.9 |
|  | **GA** | 73 | 4.9 |
|  | **AP** | 51 | 3.4 |
|  | **SP** | 33 | 2.2 |
|  | **AA** | 14 | 0.9 |
|  | **SA** | 13 | 0.9 |
|  | **AT** | 12 | 0.8 |
|  | **AV** | 9 | 0.6 |
|  | **YP** | 9 | 0.6 |
|  | **AS** | 6 | 0.4 |
|  | **GS** | 4 | 0.3 |
|  | **GV** | 3 | 0.2 |
|  | **TT** | 3 | 0.2 |
|  | **YA** | 2 | 0.1 |
|  | **GT** | 1 | <0.1 |
|  | **AY** | 1 | <0.1 |
|  | **DT** | 1 | <0.1 |
|  | **GF** | 1 | <0.1 |
|  | **GQ** | 1 | <0.1 |
|  | **NV** | 1 | <0.1 |
|  | **SN** | 1 | <0.1 |
|  | **SR** | 1 | <0.1 |
|  | **ST** | 1 | <0.1 |
|  | **SV** | 1 | <0.1 |
| Gly172 | **G** | 1389 | 92.6 |
|  | **A** | 40 | 2.7 |
|  | **S** | 36 | 2.4 |
|  | **T** | 15 | 1.0 |
|  | **E** | 6 | 0.4 |
|  | **C** | 6 | 0.4 |
|  | **D** | 1 | <0.1 |
|  | **V** | 1 | <0.1 |

**Table S2. Enzymatic activity of 17 recombinant legumains and 2 mutants**

| **Name** | **Species** | **G^237^** | **G^167^P^168^** | **G^172^** | **pH Optima** | **C/H ratio** |
| --- | --- | --- | --- | --- | --- | --- |
| VaPAL1 | *Viola albida* | I | AP | G | 6.5 | >20 **^a^** |
| VoPAL1 | *Viola orientalis* | V | AP | G | 6.5 | >20 |
| VvPAL1 | *Viola verecunda* | I | AP | G | 6.0 | >20 |
| VuPAL1 | *Viola uliginosa* | I | AP | G | 6.0 | >20 |
| VyPAL4^4^ | *Viola yedoensis* | I | AP | G | 6.0 | >20 |
| VyPAL5^4^ | *Viola yedoensis* | I | AP | G | 6.5 | >20 |
| HaPAL1 | *Helianthus annuus* | I | AA | G | 7.0 | >20 |
| BmAEP1-S161A | *Momordica charatis* | A | **A**A | G | 5.5 | >20 |
| PiAEP1 | *Psychotria ipecacuanha* | A | AP | G | 6.0 | 10.40 ± 0.56 |
| PeAEP2 | *Petunia exserta* | V | GP | G | 7.0 | 7.21 ± 0.28 |
| PeAEP1-S169G | *Petunia exserta* | A | AP | **G** | 6.5 | 3.08 ± 0.59 |
| LsAEP1 | *Lactuca sativa* | G | AP | G | 5.0 | 1.49 ± 0.30 |
| VtAEP1 | *Viola tricolor* | V | YP | G | 6.0 | 0.55 ± 0.04 |
| PeAEP1 | *Petunia exserta* | A | AP | S | 6.0 | 0.49 ± 0.10 |
| CrAEP1 | *Catharanthus roseus* | A | GP | G | 4.0 | 0.23 ± 0.02 |
| BrAEP1 | *Brassica chinensis* | G | GP | G | 5.5 | 0.17 ± 0.06 |
| BmAEP1 | *Momordica charatis* | A | SA | G | 5.5 | 0.08 ± 0.04 |
| Butelase-2 | *Clitoria ternatea* | G | GP | G | 6.0 | 0.03 ± 0.01 |
| DcAEP1 | *Dianthus caryophyllus* | G | GA | G | 6.0 | 0.02 ± 0.01 |

a: No hydrolytic product detected. b: No cyclic product detected. Highlight in red: hydrolysis-promoting variation. Bold and underlined: mutation site.
